# PP2A-B55α Terminates Mitophagy by Selectively Dephosphorylating Parkin via a Non-Canonical Interface

**DOI:** 10.64898/2026.11.23.768473

**Authors:** Fuxiang Liang, Lingsheng Meng, Lulong Huang, Shangxiang Ye, Xingyu Liu, Zefeng Nie, Xu Dong, Fangfang Li, Zhou Gong, Wei Yang, Weiping Zhang, Chun Tang

## Abstract

Mitophagy, the selective autophagic clearance of damaged mitochondria, is tightly governed by the PINK1–Parkin signaling axis. Although the activation of Parkin—driven by phosphorylation of ubiquitin (Ub) and Parkin—has been well characterized, the mechanisms that terminate mitophagy remain poorly understood. Here, we identify the heterotrimeric protein phosphatase 2A (PP2A) holoenzyme assembled with the B55α regulatory subunit (PP2A–B55α) as the terminator of mitophagy. We demonstrate that PP2A–B55α preferentially dephosphorylates pParkin over pUb in cells, effectively blocking Parkin mitochondrial translocation. Mechanistically, B55α binds Parkin through an interface distinct from the one commonly used to engage PP2A substrates containing disordered short linear motifs (SLiMs), thereby accommodating the folded Parkin ubiquitin-like (Ubl) domain. Leveraging this recognition mechanism, we rationally designed a Parkin-derived competitive inhibitor that validates this non-canonical interface and selectively uncouples pParkin dephosphorylation from the global pUb pool. Together, our findings define a negative regulatory mechanism governing Parkin and provide a proof-of-concept strategy to selectively enhance mitophagy without disrupting broader Ub signaling.

## Introduction

Mitophagy is a selective autophagic process that removes damaged mitochondria to preserve organelle quality and cellular homeostasis. Dysregulation of mitophagy is implicated in neurodegenerative conditions, including Alzheimer’s and Parkinson’s diseases (1, 2). Initiation of mitophagy is governed by the PINK1-Parkin axis. Upon mitochondrial depolarization, the kinase PINK1 is stabilized and activated on the outer mitochondrial membrane (OMM), which then phosphorylates ubiquitin (pUb) at residue S65 (3–5). pUb binding to Parkin allosterically displaces Parkin’s N-terminal ubiquitin-like (Ubl) domain, rendering Ubl accessible for PINK1 phosphorylation, also at residue S65 (6–8). This dual-phosphorylation mechanism thus unleashes Parkin’s E3 ligase activity, allowing pParkin to translocate to mitochondria and ubiquitylate OMM proteins required to execute mitophagy (7–10). As such, Parkin is the central, rate-limiting hub for mitophagy initiation.

Conversely, the negative regulatory mechanisms to prevent excessive, potentially detrimental mitochondrial clearance have been less characterized. Several deubiquitinases (DUBs), including USP14, USP15, and USP30, are known to antagonize Parkin-mediated mitophagy by removing polyubiquitin chains from OMM substrates (11–13). Furthermore, phosphatases such as PTEN-L and PPEF2 have been identified to counteract PINK1 by dephosphorylating pUb (14, 15). However, the expression of PTEN-L—a variant of the PTEN lipid phosphatase—depends on non-canonical translation initiation and is therefore highly context-dependent, whereas PPEF2 expression is largely restricted to the neural layer of the retina.

Given the ubiquitous distribution of mitochondria and the universal requirement for mitochondrial quality control across tissue types, neither PTEN-L nor PPEF2 meet the criteria of a generalized, systemic regulator of mitophagy. Furthermore, as pUb participates in a multitude of cellular activities beyond mitophagy (16–18), modulation of pParkin, instead of pUb, can be a more targeted stratagem for the cell.

Here, we report that the protein phosphatase 2A (PP2A) holoenzyme containing B55α regulatory subunit (B55α) directly dephosphorylates both pParkin and pUb, functioning as a ubiquitous brake on PINK1-initiated mitophagy. Crucially, we discover that PP2A-B55α preferentially dephosphorylates pParkin over pUb. Capitalizing on this substrate divergence, we designed a peptide that acts as a specific inhibitor against pParkin dephosphorylation. Our study delineates a fundamental molecular mechanism governing Parkin activity and provides a proof-of-principle for sustaining mitophagy without perturbing broader Ub signaling.

## Results

### PP2A core enzyme dephosphorylates both pUb and pParkin *in vitro*

Given the structural homology between Parkin Ubl domain and Ub, particularly their shared phosphorylation at residue S65, we hypothesized that they are targeted by the same ubiquitous phosphatase. Because Ub is highly amenable to isotopic labeling for high-resolution NMR studies (19), we first utilized pUb as a model substrate. Incubation of pUb with HeLa cell lysate reverted the protein to unmodified state, as evidenced by changes in two-dimensional ^1^H-^15^N heteronuclear single-quantum coherence (HSQC) NMR spectra (Fig. S1A). The appearance of new peaks was not a consequence of protein degradation, as they could be unambiguously assigned to unmodified Ub (Fig. S1B). The data thus show that the cell lysate possesses specific phosphatase activity directed against pUb.

To monitor pUb dephosphorylation as it happens, we introduced a Cys point mutation at Ub K48 site (20) and conjugated a trifluoromethyl probe, for the collection of a series of one-dimensional ^19^F-NMR spectra. Previous studies have established that pUb alternates between two conformational states—the relaxed and retracted states—which manifest as two distinct ^19^F NMR signals (8, 19, 21) (Fig. S1C). The addition of cell lysate caused the gradual disappearance of these two peaks, with the concomitant appearance of a new downfield peak corresponding to the unmodified Ub (Fig. S1C). To identify the specific phosphatase responsible, we conducted a screening assay using class-specific small-molecule inhibitors. Treatment with either okadaic acid or cantharidin significantly suppressed the phosphatase activity in the cell lysate and slowed pUb dephosphorylation (Fig. S1D-H). Because okadaic acid and cantharidin are specific inhibitors of the PP2A and PP1 families, respectively (22, 23), this ^19^F-NMR-based screening approach narrowed our primary pUb phosphatase candidates to these two enzymes.

We subsequently prepared recombinant PP2A and PP1 core enzymes and titrated them into the ^19^F-labeled pUb. The addition of 0.5 µM PP2A rapidly eliminated the two pUb peaks and accelerated the appearance of the unmodified Ub peak. In contrast, the addition of 0.5 µM PP1 core enzyme to an equivalent amount of pUb induced only small changes in peak intensities compared to the untreated control (Fig. S2A, B). Further assessment using HSQC NMR confirmed the rapid and complete conversion of PP2A-treated pUb to unmodified Ub (Fig. 1A). Collectively, the results demonstrate that PP2A is a *bona fide* pUb phosphatase.

**Fig. 1.**
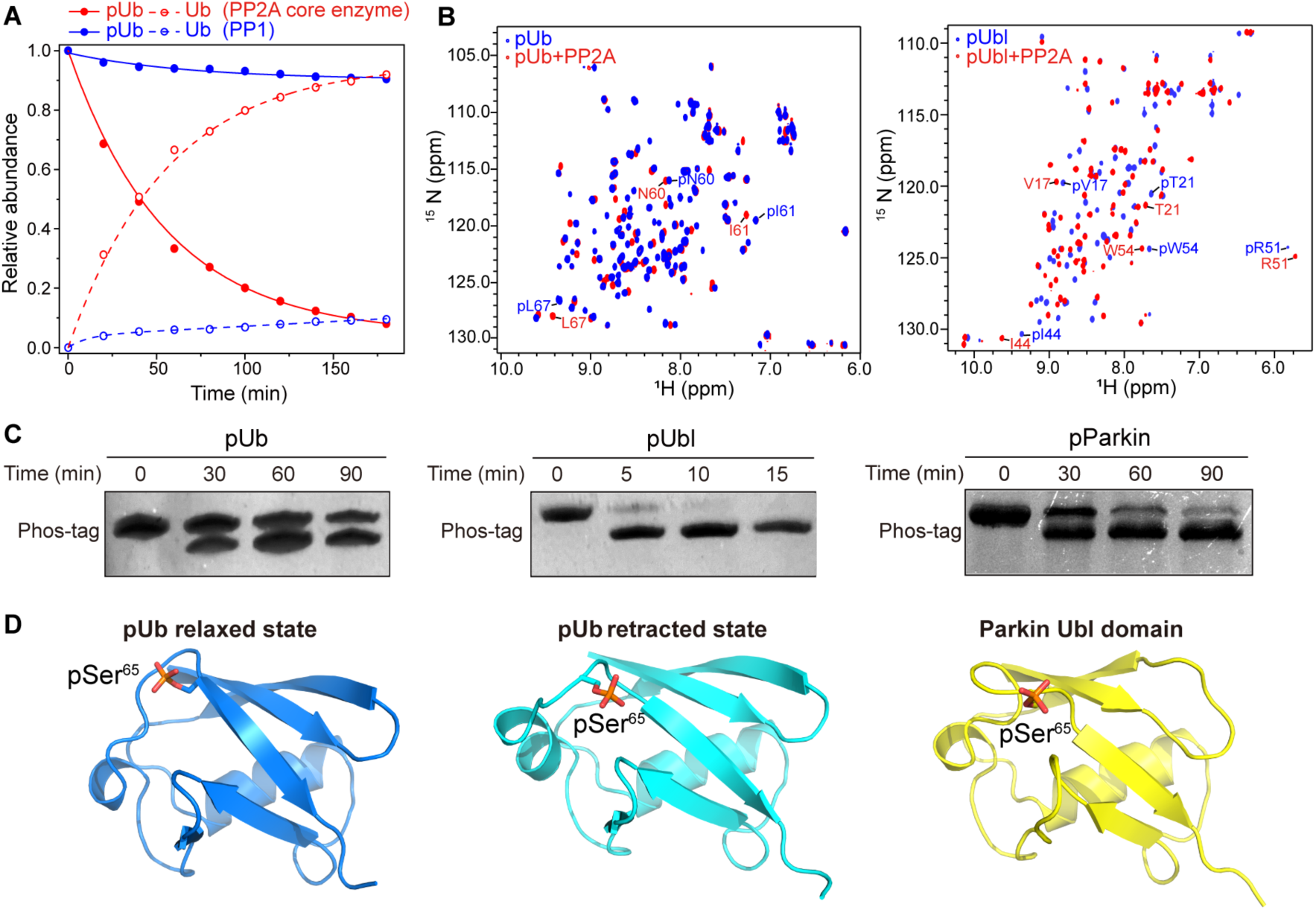
PP2A core enzyme directly dephosphorylates pUb and pParkin *in vitro*. **(A)** Dephosphorylation kinetics of pUb treated with PP2A or PP1 core enzyme. Data were normalized to the initial ^19^F peak intensities. The unmodified Ub peak intensity was calculated by subtracting the remaining pUb intensity from the total intensity. **(B)** Overlay of 2D ^1^H-^15^N HSQC spectra of 100 µM pUb (left) and Parkin pUbl (residues 1-76, right) before and after 1 h incubation with 0.5 µM PP2A core enzyme. Well-isolated peaks are labeled. **(C)** Phos-tag gel analysis comparing the dephosphorylation rates of pUb (left), isolated Parkin pUbl domain (middle), and full-length PARKIN (right) by PP2A core enzyme. Representative gels from *n* = 3 independent experiments are shown. **(D)** Structural representations of pUb relaxed state (left, PDB: 5XK5), pUb retracted state (middle, PDB: 5XK4), and Parkin pUbl domain (right, PDB: 5TR5). The structures illustrate higher accessibility of pSer65 in Parkin Ubl and pUb retracted state than in pUb relaxed state.

Having validated pUb as a substrate, we next investigated whether PP2A core enzyme could also dephosphorylate pParkin. We observed that PP2A robustly dephosphorylates the isolated phosphorylated Ubl domain (pUbl) of PARKIN, as the addition of 0.5 µM PP2A core enzyme completely restored the NMR spectrum to that of unphosphorylated Ubl (Fig. 1B). Notably, PP2A dephosphorylated Parkin pUbl domain with markedly accelerated kinetics compared to pUb. Phos-tag gel analysis revealed that the pUbl band disappeared within 15 min of PP2A addition, whereas the pUb band intensity was only reduced by ∼50% after 30 min (Fig. 1C). This rapid processing was similarly observed for full-length pParkin, albeit at a slightly reduced rate, which can be attributed to steric hindrance of the Ubl domain within the full-length protein (Fig. 1D).

This inherent difference in kinetics can be rationalized structurally. In Parkin, pSer^65^ is situated in an extended, highly accessible loop, whereas in Ub, pSer^65^ only becomes fully exposed in the retracted state (Fig. 1E) (8, 19). Indeed, PP2A dephosphorylated T66V/L67N double-mutant of Ub, engineered to constitutively adopt the retracted state (24), much more efficiently than the wild-type pUb (Fig. S2C, D). Thus, we establish that the PP2A core enzyme dephosphorylated both pUb and pParkin, and that the basal catalytic efficiency is dictated by the conformational accessibility of the phosphoryl group.

### B55α is the regulatory subunit for PP2A-mediated Parkin dephosphorylation

A fully functional PP2A holoenzyme is a heterotrimer comprising a scaffolding subunit (A), a catalytic subunit (C), and a regulatory subunit (B) (25–27). The highly variable B subunit plays a crucial role in controlling PP2A activity by dictating substrate selectivity and directing the subcellular localization of the holoenzyme (28–31). To identify the specific regulatory subunit responsible for mitophagy regulation, we used an interactome screening strategy. We reasoned that during mitophagy, the massive accumulation of pUb chains on the OMM forms a highly concentrated signaling platform, which could physically recruit the mitochondrial phosphatase complex to proximity. We thus utilized FLAG-tagged pUb as a bait to capture the endogenous phosphatase holoenzyme. Specifically, we incubated cell lysates with recombinant FLAG-tagged pUb and the GST-tagged PP2A scaffolding subunit, performed mild chemical cross-linking to preserve transient interactions, and isolated the interacting proteins using consecutive FLAG and GST affinity purifications (Fig. 2A). Through this dual-affinity approach coupled with high-resolution liquid chromatography-tandem mass spectrometry (LC-MS/MS), we identified multiple peptides derived from the regulatory subunit B55α in the eluted mixture (Fig. S3A-C).

**Fig. 2.**
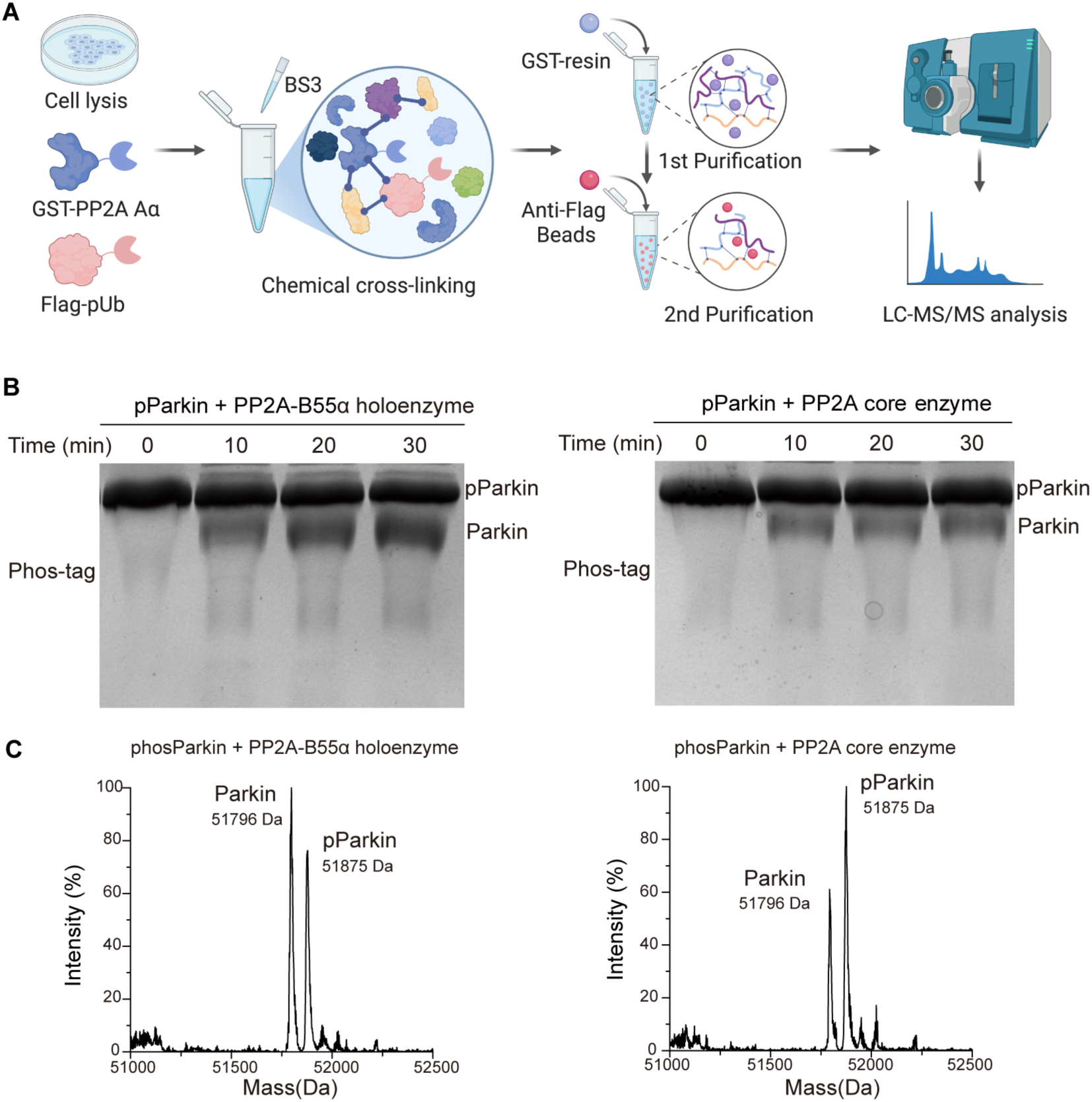
B55α is the regulatory subunit for PP2A-mediated pParkin dephosphorylation. (**A**) Schematic illustration of the dual-affinity interactome screening workflow. HeLa cell lysates were incubated with recombinant GST-tagged PP2A scaffolding subunit Aα and FLAG-tagged pUb, followed by mild chemical cross-linking with the cross-linker BS_3_. The assembled complexes were sequentially purified using GST-resin and anti-FLAG beads, and the final eluates were subjected to LC-MS/MS analysis to identify the regulatory subunit. (**B**) Phos-tag SDS-PAGE analysis comparing the dephosphorylation kinetics of full-length pParkin treated with reconstituted PP2A– B55α holoenzyme (left) versus PP2A core enzyme alone (right) over time (0–30 min). Dephosphorylated Parkin exhibits faster electrophoretic mobility than mobility-retarded pParkin; the dephosphorylated band displays consistently higher intensities across time points when catalyzed by the holoenzyme. (**C**) Intact protein LC-MS spectra of pParkin following dephosphorylation by equimolar amounts of PP2A holoenzyme (left) or core enzyme (right) after 40 min of incubation at room temperature. The observed mass shift of -79 Da matches the loss of a single phosphoryl group (51,796 Da for Parkin vs. 51,875 Da for pParkin). Theoretical molecular weights (51,795 Da for Parkin and 51,874 Da for pParkin) account for two extra N-terminal residues (Pro-Gly) retained following PreScission protease cleavage of the GST tag. Inclusion of the B55α subunit in the holoenzyme increases the ratio of dephosphorylated Parkin over pParkin from ∼58% to ∼119%.

To validate whether B55α functions as the authentic regulatory subunit for pParkin and pUb dephosphorylation, we recombinantly purified B55α and reconstituted the PP2A holoenzyme. This was achieved by combining the B55α subunit with the pre-assembled heterodimeric core enzyme (composed of the catalytic C subunit and the PR65α scaffolding subunit) at equal molar amount. The inclusion of the B55α subunit accelerated pUb dephosphorylation, as quantified by real-time ^19^F-NMR (Fig. S3D, E). Crucially, the reconstituted PP2A-B55α holoenzyme catalyzed the dephosphorylation of full-length pParkin much more efficiently than the core enzyme alone, as evidenced by time-course Phos-tag gel analysis (Fig. 2B). This B55α-mediated kinetic acceleration was further corroborated by intact mass spectrometry analysis; treatment with the PP2A holoenzyme resulted in a rapid, near-complete mass shift to the value corresponding to the unmodified Parkin, about twice as much faster than observed with the PP2A core enzyme alone (Fig. 2C). Together, these results demonstrate that B55α associates with the PP2A core enzyme to form a highly active, functional holoenzyme capable of efficiently dephosphorylating full-length pParkin.

### B55α dictates the specific dephosphorylation of pParkin in cells

Administration of the mitochondrial uncoupler carbonyl cyanide chlorophenyl hydrazone (CCCP) or the respiratory chain inhibitors oligomycin and antimycin A (O/A) dissipates the mitochondrial membrane potential, thereby triggering PINK1-Parkin-mediated mitophagy (32, 33). To investigate how PP2A counteracts PINK1 kinase activity in cells, we modulated the protein level of the PP2A core enzyme, i.e., the catalytic subunit (Cα). Transient overexpression of Cα markedly attenuated the accumulation of both pParkin and pUb that was otherwise strongly elicited by CCCP or O/A treatment (Fig. 3A). In contrast, overexpression of the PP1 catalytic subunit α (PP1α) did not significantly alter CCCP-induced pParkin or pUb levels, supporting a differential effect of these two phosphatases under the conditions tested (Fig.3B). Conversely, siRNA-mediated knockdown of PP2A Cα exacerbated the accumulation of pParkin and pUb, demonstrating an additive effect with the mitophagy inducers on substrates that would otherwise be rapidly hydrolyzed (Fig. 3C). Importantly, the protein levels of total Parkin and Ub remained unaffected regardless of PP2A Cα expression levels, ruling out protein degradation as a confounding factor. The data thus indicate that the PP2A core enzyme dephosphorylates both pParkin and pUb in the cellular environment.

**Fig. 3.**
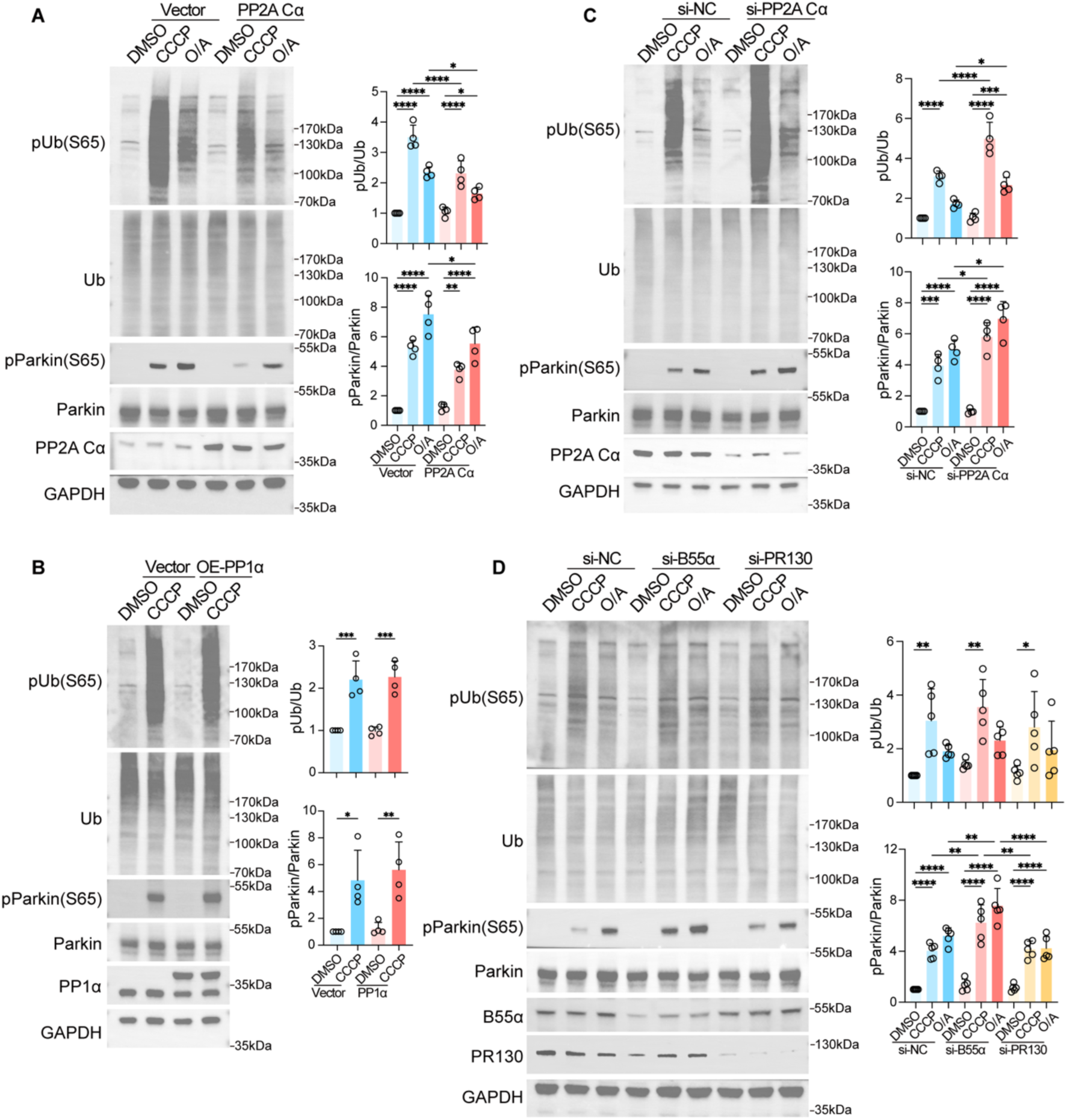
B55α dictates the specific dephosphorylation of pParkin in cells. (**A**) Overexpression of the PP2A catalytic subunit (Cα) decreases pParkin and pUb levels. HeLa cells were transiently transfected with an empty vector or a plasmid expressing PP2A Cα, followed by treatment with DMSO, CCCP (10 µM for 4 h), or oligomycin/antimycin A (O/A, both at 250 nM for 2 h). (**B**) Overexpression of PP1α does not significantly alter CCCP-induced pParkin or pUb levels. HeLa cells were transiently transfected with an empty vector or a plasmid expressing PP1α, followed by treatment with DMSO or CCCP (10 µM for 4 h). (**C**) Knockdown of PP2A Cα increases pParkin and pUb levels. HeLa cells were transfected with non-targeting control siRNA (si-NC) or siRNA specific for PP2A Cα, followed by the indicated treatments. (**D**) Knockdown of B55α selectively increases pParkin levels without significantly altering pUb levels, whereas knockdown of PPP2R3A, another PP2A regulatory subunit, does not significantly alter either pParkin or pUb levels. HeLa cells were transfected with si-NC or siRNA targeting B55α (si-B55α) or PPP2R3A (si-PR130), followed by the indicated treatments. For (A–D), representative immunoblots and densitometric quantification of pUb/Ub and pParkin/Parkin ratios are shown. GAPDH served as a loading control. Data are presented as mean ± SEM from n = 4-5 independent experiments. * *P* < 0.05, ** *P* < 0.01, *** *P* < 0.001, and **** *P* < 0.0001, determined by one-way ANOVA.

To determine whether the regulatory subunit B55α specifically dictates this dephosphorylation process, we evaluated the effect of B55α depletion on substrate phosphorylation. In HeLa cells treated with CCCP or O/A, Western blot analysis revealed that siRNA-mediated knockdown of B55α significantly increased the accumulation of pParkin.

Unexpectedly, B55α depletion had a negligible effect on pUb levels (Fig. 3D). A second independent siRNA targeting B55α also increased pParkin levels without significantly altering pUb levels, supporting the reproducibility of this phosphorylation pattern across different siRNA sequences (Fig. S3F). To assess regulatory-subunit specificity, we depleted another PP2A regulatory subunit, PPP2R3A, with siRNA. Unlike B55α depletion, PPP2R3A knockdown did not significantly alter pParkin or pUb levels following CCCP or O/A treatment (Fig.3D). Though appearing discordant with our *in vitro* observations (Fig. 2 and Fig. S3), this discrepancy likely reflects the distinct structural features and the physiological redundancy of pUb and pParkin substrates in cells. The highly accessible pUb chains on the outer mitochondrial membrane (OMM) may be redundantly targeted by multiple PP2A holoenzymes harboring different regulatory subunits, as well as by other cellular phosphatases such as PTEN-L (15). In contrast, the dephosphorylation of pParkin requires the precise scaffolding, substrate specificity, and spatial positioning provided by the B55α subunit. Taken together, while the catalytic core of PP2A is capable of dephosphorylating both substrates, B55α functions as the regulatory subunit enabling a preference for pParkin.

### PP2A-B55α holoenzyme suppresses Parkin mitochondrial translocation and mitophagy

Parkin translocation to depolarized mitochondria is a prerequisite for the initiation of mitophagy, where, functioning as an E3 ligase, it ubiquitinates a multitude of downstream OMM proteins (10, 34). To investigate whether PP2A-mediated dephosphorylation affects this translocation process, we utilized HeLa cells stably expressing YFP-tagged Parkin (10). Upon treatment with CCCP or O/A, YFP-Parkin was robustly recruited to mitochondria, as evidenced by a progressive increase in colocalization with the mitochondrial protein TOM20 (Fig. S4A). As expected, transient overexpression of the PP2A catalytic subunit (Cα) significantly attenuated Parkin mitochondrial translocation across all tested time points (Fig. S4A).

To confirm this inhibitory effect at the single-cell level and to exclude potential transfection artifacts, we introduced a C-terminally mCherry-tagged PP2A Cα construct (PP2A-Cα-mCherry) into the cell line. Under basal conditions, PP2A-Cα-mCherry was diffusely distributed throughout the cytoplasm and did not alter baseline Parkin localization (Fig. 4A). Following CCCP or O/A treatment, robust YFP-Parkin mitochondrial translocation was exclusively observed in untransfected (mCherry-negative) cells. In contrast, adjacent cells expressing PP2A-Cα-mCherry exhibited a near-complete blockade of YFP-Parkin recruitment to the mitochondria (Fig. 4A, indicated by white arrows).

**Fig. 4.**
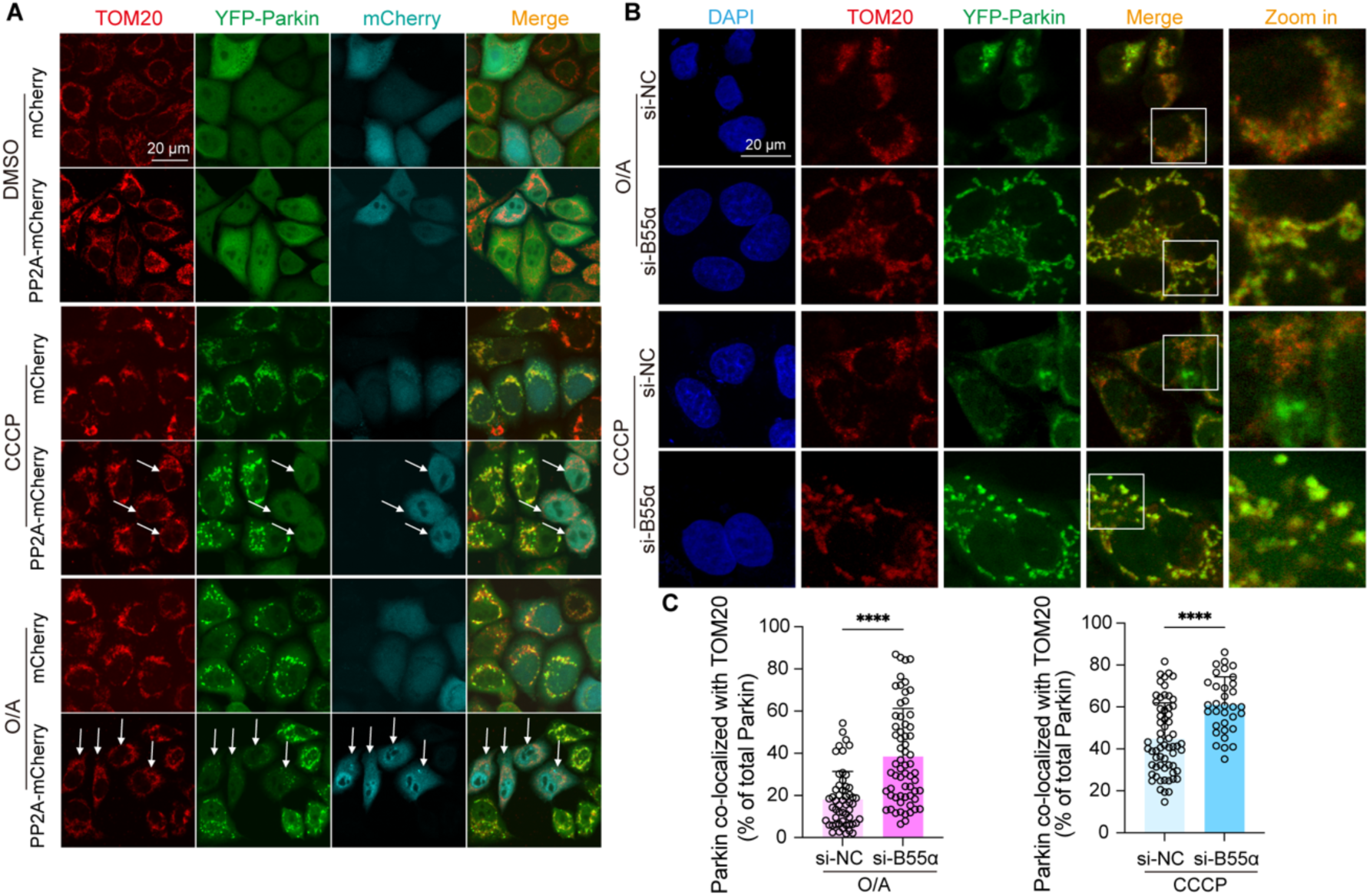
The PP2A-B55α holoenzyme inhibits Parkin mitochondrial translocation. (**A**) Overexpression of PP2A Cα blocks Parkin translocation at the single-cell level. HeLa cells stably expressing YFP-Parkin (green) were transiently transfected with mCherry (Vector control) or PP2A-Cα-mCherry. Cells were treated with DMSO, CCCP (10 µM for 2 h), or oligomycin/antimycin A (O/A, both at 250 nM for 1 h). Note that mCherry fluorescence is pseudo-colored cyan to distinguish it from the TOM20 staining (red). Arrows indicate PP2A-Cα-mCherry-expressing cells that fail to recruit YFP-Parkin to mitochondria. (**B**) Knockdown of the regulatory subunit B55α enhances Parkin mitochondrial translocation. HeLa cells stably expressing YFP-Parkin were transfected with si-NC or siRNA specific for B55α, followed by the indicated treatments. Mitochondria were delineated by TOM20 immunofluorescence (red). (**C**) Quantitative analysis of Parkin colocalization with TOM20 from the experiments in (B). Data are presented as mean ± SEM from *n* = 3 independent experiments (at least 15 cells were analyzed per condition). **** *P* < 0.0001, determined by unpaired Student’s *t*-test.

Having established that PP2A suppresses Parkin translocation, we next evaluated its impact on downstream mitophagic flux using the Mito-QC reporter system (35). In this assay, the delivery of mitochondria to acidic lysosomes is visualized as a fluorescent shift from dual green/red to solely red puncta. Consistent with the translocation data, overexpression of PP2A Cα significantly suppressed the formation of mito-lysosomes induced by CCCP or O/A (Fig. S4C). Conversely, siRNA-mediated depletion of PP2A Cα markedly enhanced the accumulation of mito-lysosomes under identical stress conditions (Fig. S4C). These results indicate that PP2A-mediated dephosphorylation of pUb and pParkin effectively attenuates the progression of mitophagy.

To substantiate the physiological requirement for the regulatory subunit B55α in suppressing Parkin mitochondrial translocation, we examined the effect of B55α depletion. In line with our biochemical observation of pParkin accumulation (Fig. 3D), immunofluorescence imaging revealed that B55α knockdown significantly enhanced the colocalization of YFP-Parkin with TOM20 following CCCP or O/A treatment (Fig. 4B, C). To assess whether the effects of PP2A Cα or B55α depletion were associated with altered mitochondrial membrane potential, we performed JC-1 staining. CCCP significantly reduced the JC-1 red/green fluorescence ratio in all three siRNA groups. However, no significant differences in this ratio were detected among si-NC-, si-PP2A Cα-, and si-B55α-transfected cells under either DMSO- or CCCP-treated condition (Fig. S4B). Together, these imaging and functional results establish that the PP2A-B55α holoenzyme specifically dephosphorylates pParkin, preventing unnecessary mitophagy by restricting Parkin mitochondrial translocation and subsequent activation.

### Structural modeling and validation of PP2A-B55α-Parkin interaction

We utilized AlphaFold (36) to model the complex structure between the heterotrimeric PP2A-B55α holoenzyme and the Parkin Ubl domain (Fig. 5A). The predicted model reveals a bipartite interaction mode that coordinates the dephosphorylation event, with the productive pose recovered across ∼60% of the models of independent seeded runs (Fig. S5A). On one hand, pSer^65^ in the folded Parkin Ubl domain docks onto PP2A catalytic subunit Cα in close proximity to its binuclear metal center, consistent with the desired chemistry (Fig. 5B). On the other hand, the Ubl domain engages the regulatory subunit B55α to confer substrate specificity. Inter-subunit Predicted Aligned Error (PAE) analysis (Fig. S5B) identified a cluster of acidic residues in B55α (E91, E93, and E94) that participates in electrostatic interactions with basic residues R72 and R75 flanking the C-terminus of the Parkin Ubl domain (Fig. 5B).

**Fig. 5.**
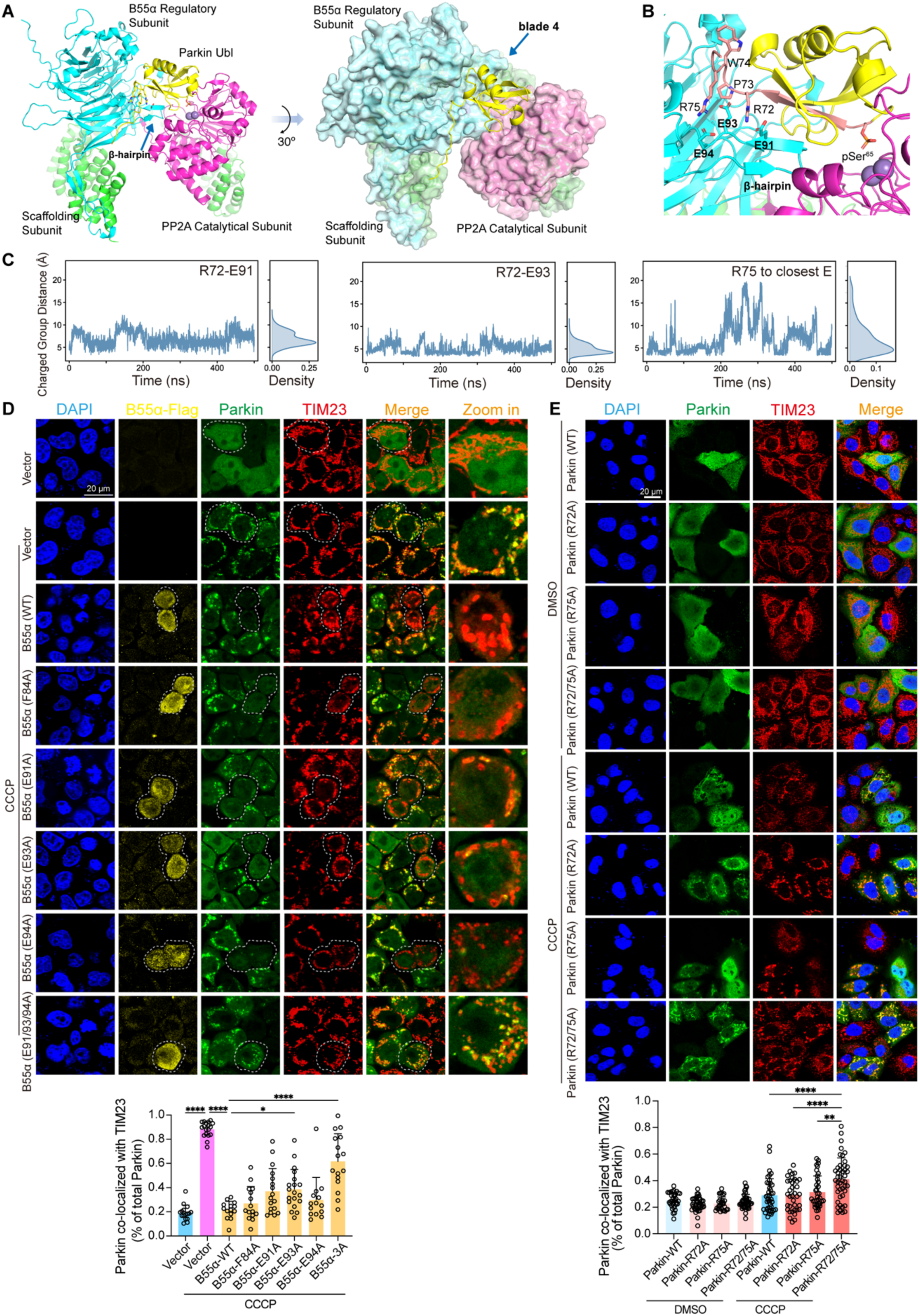
Structural basis of the B55α-Parkin interaction and mutational validation. **(A)** Overall structural model of the PP2A-B55α-Parkin Ubl complex predicted by AlphaFold. The heterotrimeric PP2A holoenzyme consists of the scaffolding subunit (green), the regulatory subunit B55α (cyan), and the catalytic subunit (magenta). The Parkin Ubl domain is shown in yellow. Surface representation on the right highlights the canonical interface of top groove in B55α is away from the current interface engaging Parkin Ubl domain. **(B)** Detailed view of the regulatory and catalytic interfaces. The phosphorylated S65 residue of Parkin Ubl (yellow) docks into the catalytic pocket. Key interacting residues between the Parkin Ubl C-terminal tail (R72, P73, W74, R75) and the acidic patch (E91, E93, and E94) on the B55α β-hairpin (cyan) are indicated. **(C)** All-atom molecular dynamics (MD) simulations of the PP2A-B55α-Parkin Ubl complex. The plots show the distances between the charged groups of Parkin R72 and B55α E91 (left), Parkin R72 and B55α E93 (middle), and Parkin R75 to the closest glutamate residue on B55α (right) over time (ns), alongside their corresponding density distributions. **(D)** Effects of B55α interface mutations on Parkin mitochondrial translocation. HeLa cells stably expressing Parkin (green) were transiently transfected with an empty vector or Flag-tagged wild-type B55α or the indicated mutants (yellow). Cells were treated with CCCP (10 µM for 2 h) as indicated; a vector-transfected control without CCCP treatment is also shown. B55α-3A denotes the E91A/E93A/E94A triple mutant. Mitochondria were immunostained for TIM23 (red). Nuclei were counterstained with DAPI (blue). White dashed lines outline the cells of interest. Enlarged views are shown in the rightmost column. The graph below quantifies the fraction of total Parkin signal colocalized with TIM23. **(E)** Validation of interfacial residues in Parkin Ubl domain. Representative confocal immunofluorescence images of HeLa cells expressing wild-type Parkin, R72A or R75A single mutants, or R72A/R75A double mutant (green) treated with vehicle (DMSO) or CCCP (10 µM, 1 h). Mitochondria were counterstained with anti-TIM23 (red), and nuclei were stained with DAPI (blue). Scale bar, 20 µm. Lower panels: Quantitative analysis of Parkin recruitment to depolarized mitochondria, plotted as the percentage of total Parkin signal colocalized with TIM23. Data represent mean ±SEM from *n* = 3 independent biological replicates (>15 cells analyzed per condition; * *P* < 0.05, ** *P* < 0.01, and **** *P* < 0.0001, one-way ANOVA followed by Tukey’s multiple comparisons test).

Under the canonical model of PP2A substrate recruitment, substrate specificity is driven by a seven-repeat β-propeller architecture in B55α, wherein a short linear motif (SLiM) within the intrinsically disordered region (IDR) of a substrate protein docks to a conserved acidic groove of B55α β-propeller structure, centered around residue D197 in blade 4 (37–39). In the case of Parkin, it is not possible for the rigid, folded Ubl domain to engage this top groove interface while at the same time, position the phosphoryl group at the active site of the catalytic subunit.

Consequently, the Ubl domain has to dock “lower”, with the B55α interface comprising a cluster of negatively charged residues in a protruding β-hairpin (Fig. 5A, B). This non-canonical interaction thus resolves the distance constraint for a folded PP2A substrate protein.

This novel interaction mode was further corroborated by all-atom molecular dynamics (MD) simulations. Over trajectories spanning hundreds of nanoseconds, the PP2A-B55α-Parkin assembly remained highly stable. Specifically, we monitored the distances between the key charged groups at the interface. The distances between the guanidine group of Parkin R72 and the carboxyl groups of B55α E91 and E93, as well as the distance from Parkin R75 to its closest glutamate residue of B55α, exhibited significant retention at the interface (Fig. 5C). The density distributions confirm that these electrostatic contacts form mostly stable interactions, ensuring the Ubl domain anchored at B55α.

To validate the functional relevance of the predicted B55α-Parkin interface, we mutated the key acidic residues in B55α and assessed their ability to rescue the PP2A-mediated inhibition of mitophagy in cells. In agreement with the role of PP2A holoenzyme as a negative regulator of mitochondrial clearance, transient overexpression of wild-type B55α (B55α-WT) robustly blocked the translocation of Parkin to TIM23-positive damaged mitochondria upon CCCP treatment, whereas adjacent untransfected cells in the same field of view displayed pronounced Parkin recruitment (Fig. 5D). In comparison, mutating the acidic residues reversed this inhibitory effect, rescuing Parkin translocation despite the high-level expression of B55α. The triple-alanine mutant B55α-3A (E91A/E93A/E94A), which largely neutralizes this acidic interface, exhibited the most pronounced rescue effect of Parkin colocalization with TIM23 (Fig. 5D). Mutation of residue F84, also located in the β-hairpin, however, showed a much weaker reversal trend (Fig. 5D).

Complementary to the B55α mutational analysis, we examined the reciprocal basic residues in the Parkin Ubl domain by generating the R72A and R75A single mutants and the R72A/R75A double mutant. Under DMSO-treated conditions, wild-type Parkin and the mutants showed broadly similar subcellular distributions. Following CCCP treatment, the R72A/R75A double mutant exhibited significantly greater colocalization with TIM23 than wild-type Parkin or either single mutant (Fig. 5E). These findings corroborate the B55α mutational results and confirm the predicted interface for Parkin recruitment to PP2A holoenzyme.

Together, these structural, computational, and functional findings established a non-canonical binding mode for the Parkin Ubl domain to the B55α regulatory subunit, providing a compelling explanation for the specific PP2A-B55α-mediated termination of mitophagy.

### Competitive interference of Parkin dephosphorylation enhances mitophagy

Having mapped the structural features of Parkin-B55α interface, we hypothesized that disrupting this non-canonical interaction would be a targeted strategy to sustain Parkin phosphorylation and enhance mitophagy. Upon comparing the primary sequence of the Parkin Ubl domain with that of Ub, several key structural differences emerge. While the Ub C-terminal tail (residues 70–76: VLRLRGG) terminates in a neutral, flexible diglycine motif (Gly^75^–Gly^76^), the Parkin Ubl tail carries a higher positive charge density owing to the basic residues Arg^75^–Lys^76^. Furthermore, the spacing of the positively charged arginine residues differs between the two tails: the two arginine residues in the Parkin Ubl tail (residues 69–76: VQRPWRKG) are separated by two residues (Pro^73^ and Trp^74^), whereas the corresponding residues in the Ub tail are separated by a single leucine residue (Leu^72^ in Ub alignment).

To systematically interrogate this interface and to explore structure-activity relationships, we synthesized a series of FITC-conjugated short peptides derived from the Parkin-Ubl C-terminus. This panel included a wild-type (WT) mimetic peptide corresponding to residues 71–77 of the Parkin Ubl domain (PP#1) and three variants (PP#2–4) harboring single amino acid substitutions at the conserved Trp^74^ residue (substituted with Tyr, Leu, or Gly, where W74L mimics the hydrophobic sequence of the Ub tail) (Fig. 6A). Physical interaction was confirmed by fluorescence polarization (FP) *in vitro* for PP#1 with the titration of a disulfide-stabilized B55α β-hairpin peptide (Fig. S6A). Owing to the absence of cooperative tertiary contacts and electrostatic steering provided by the intact protein, it is expected that the isolated peptide fragments display moderate binding affinity (Fig. S6B). Yet the data confirm direct physical interaction of the peptide with the B55α recognition motif.

**Fig. 6.**
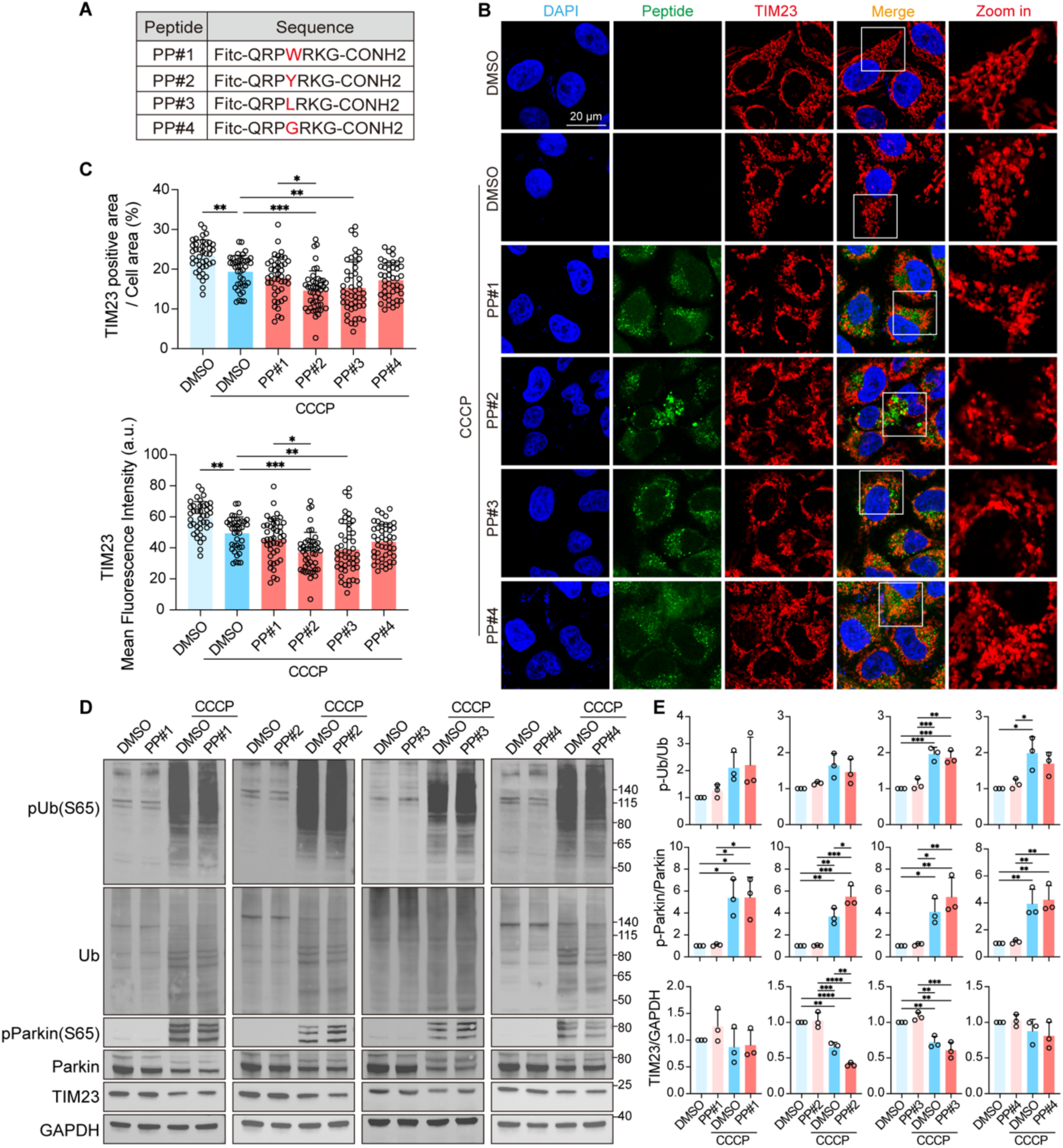
Competitive interference of Parkin dephosphorylation enhances mitophagy. (**A**) Sequences of the synthesized FITC-labeled peptides derived from the Parkin Ubl tail. PP#1 represents the wild-type sequence, while PP#2–4 contain single amino acid substitutions. (**B**) Representative immunofluorescence images showing the effect of peptide treatments on mitochondrial clearance. HeLa cells were pre-treated with the indicated peptides (10 µM for 4 h) or DMSO, followed by CCCP treatment (10 µM for 2 h). The intracellular distribution of the peptides is indicated by the FITC signal (green), and mitochondria were immunostained for TIM23 (red). White boxes indicate regions shown in the magnified views. (**C**) Quantitative analysis of the TIM23-positive area per cell area (%) and the mean fluorescence intensity of TIM23 from (B). Data are presented as mean ± SEM from *n* = 3 independent experiments (>15 cells analyzed per group). (**D**) Representative Western blot analysis evaluating the specific impact of the peptides on the phosphorylation of Parkin and Ub. Cells were treated with DMSO or the indicated peptides under basal or CCCP-induced conditions. (**E**) Densitometric quantification of p-Ub(S65), p-Parkin(S65), and TIM23 protein levels from (D). Phosphorylated protein levels were normalized to their respective total protein levels, and TIM23 was normalized to GAPDH. Data are presented as mean ± SEM from *n* = 3 independent experiments. * *P* < 0.05, ** *P* < 0.01, *** *P* < 0.001, and **** *P* < 0.0001, determined by one-way ANOVA.

Under basal, non-stressed conditions, the intracellular delivery of these peptides did not alter mitochondrial morphology or baseline mitochondrial area (Fig. S7). Upon the induction of mitophagy using the mitochondrial uncoupler CCCP, cells treated with either the wild-type mimetic (PP#1) or variants PP#3 (W74L) and PP#4 (W74G) showed no significant difference in mitochondrial clearance compared to the vehicle control. Strikingly, treatment with the tyrosine-substituted variant PP#2 (W74Y) significantly accelerated mitophagy, as demonstrated by a substantial reduction in both the TIM23-positive mitochondrial area and the mean fluorescence intensity (Fig. 6B, C). We reasoned that replacing the bulky, hydrophobic tryptophan indole ring with a tyrosine residue in PP#2 likely confers higher aqueous solubility. The hydroxyl group of the Tyr may also be well-positioned for additional hydrogen-bond with adjacent residues in B55α, thereby enabling PP#2 to outcompete endogenous full-length Parkin for holoenzyme binding.

To biochemically validate that this enhanced mitophagic flux is driven by the selective prevention of Parkin dephosphorylation, we evaluated the phosphorylation status of the core mitophagy components. Quantitative Western blot analysis revealed a highly specific, target-selective effect: upon CCCP treatment, only the PP#2 peptide sustained significantly elevated levels of pParkin relative to the CCCP-only control, whereas pUb levels remained unaffected across all peptide treatments (Fig. 6D, E). This selective preservation of pParkin, but not pUb, is in perfect agreement with our B55α knockdown data (Fig. 3D). Together, these findings demonstrate that sequence optimization of the Ubl C-terminal tail can yield an inhibitor that competitively blocks the Parkin-B55α interaction, thereby accelerating mitochondrial clearance without perturbing the global pUb pool.

## Discussion

While the initiation of mitophagy *via* the PINK1-Parkin axis has been well-established, terminating mechanisms that prevent excessive, potentially detrimental mitochondrial clearance are only beginning to be elucidated. Parkin is the central hub in the mitophagic signaling cascade, and therefore, both its activation, *via* pUb binding and Ubl domain phosphorylation, and its deactivation require exquisite regulation. In the current study, utilizing a suite of biophysical, biochemical, and cell biology approaches, we demonstrate that the heterotrimeric protein phosphatase 2A (PP2A), specifically when incorporating the B55α regulatory subunit, functions as the physiological, and likely ubiquitous, phosphatase for both pParkin and pUb. Crucially, pParkin is the preferred substrate of PP2A-B55α, and the preferential dephosphorylation blocks Parkin mitochondrial translocation and halts mitophagic progression. Notably, our findings echo a recent report that PP2A-B55α modulates multiple stages of mitophagy (40); for instance, upon initial mitochondrial insult, ULK1 is rapidly released from PP2A-B55α-mediated inhibition to prime early autophagy machinery and phagophore formation. Our finding here complements this picture by positioning the same phosphatase complex as the critical terminator of Parkin activation.

From a structural perspective, the specific dephosphorylation of Parkin by PP2A-B55α unveils a non-canonical substrate-binding paradigm. It has been established that, the B55α regulatory subunit recognizes short linear motifs (SLiMs) within intrinsically disordered regions (IDRs) of substrate proteins, typically featuring a p[ST]-P-x(4,10)-[RK]-V-x-x-[VI]-R consensus sequence (37–39). These IDRs dock onto the conserved top groove of the β-propeller structure in B55α for critical electrostatic interactions. In contrast, the Parkin Ubl domain is a well-folded, structured entity, and the primary sequence immediately following pSer^65^ deviates significantly from the established SLiM consensus sequence, which strongly suggests a distinct B55α recruitment mechanism. Indeed, distance constraints prevent the folded Ubl domain from engaging residues in blade 4 of the β-propeller of the regulatory subunit, while simultaneously inserting pSer^65^ into the active site of PP2A catalytic subunit. Our structural modeling, computational analysis, and functional validation reveal that the C-terminal tail of Parkin Ubl domain docks onto the protruding β-hairpin structure at the “lower” edge of B55α propeller, thus enabling the correct position of the phosphoserine near the active site. Interestingly, a recent cryo-EM structure of PP2A-B55α in complex with the specific inhibitor IER5 demonstrates a conceptually similar binding mode (41). In that structure, IER5 not only utilizes the canonical top-groove interface but also employs its basic residues in a protruding loop, specifically R25, to form electrostatic interactions with residues E91 and E93 in B55α (Fig. S8).

Though Parkin Ubl domain is structurally homologous to Ub, the detailed sequence divergence accounts for Parkin’s preferred association with B55α. Moreover, Parkin pSer^65^ is constantly exposed, whereas Ub pSer^65^ protrudes only when the protein exists in the retracted state (Fig. 1D). The structural and kinetic gating mechanism reconciles the apparent discrepancy between the multi-substrate activity of PP2A-B55α *in vitro* and its highly selective preference in cells—while our reconstituted system demonstrates that PP2A-B55α can accelerate the dephosphorylation of both pUb and pParkin *in vitro*, B55α depletion in cells selectively impairs pParkin dephosphorylation without altering global pUb levels. Such substrate differentiation provides a unique therapeutic window to selectively target pParkin.

The direct regulation of pParkin identified here does not exclude contributions from other phosphorylation-dependent regulators of mitophagy. Parkin-mediated mitochondrial ubiquitylation promotes TBK1 activation and adaptor recruitment, and therefore, changes in Parkin activity can influence downstream TBK1–OPTN signaling (42). However, such signaling cascade does not fully resolve whether TBK1 activation or specific OPTN phosphorylation sites are unaffected following B55α manipulation in all cellular contexts. On the other hand, receptor-mediated mitophagy pathways (e.g., NIX/BNIP3) directly engage the autophagic machinery in response to specific physiological cues without utilizing a UBL domain (42, 43). This distinction should not be equated with insensitivity to B55α manipulation, since receptor-mediated mitophagy is itself subject to phosphorylation-dependent tuning. For instance, ULK1 phosphorylates BNIP3 at Ser17 and BNIP3L/NIX at Ser35 *in vitro*, and ULK1-dependent BNIP3 phosphorylation is known to promote LC3 binding and mitophagy (44). Thus, while our study delineates a specific mechanism for Parkin dephosphorylation, the full spectrum of cellular outcomes following B55α perturbation could be broader and multifaceted.

Capitalizing on the depiction of this non-canonical Parkin-B55α interface, we developed a competitive peptide capable of modulating their interaction. A single amino acid substitution in a peptide derived from the tail of Parkin Ubl domain allows the peptide to function as an effective competitive inhibitor, blocking the recruitment of pParkin to PP2A-B55α. Crucially, this engineered peptide selectively preserves the phosphorylation of Parkin without perturbing the dephosphorylation of pUb. This specificity is of profound physiological importance. PINK1-phosphorylated ubiquitin participates in a multitude of cellular events beyond mitophagy (16, 17, 45); we have recently demonstrated that excessive accumulation of pUb can disrupt ubiquitin-proteasome system (UPS)-mediated proteostasis, contributing to the pathogenesis of neurodegeneration (16). Therefore, by deploying a peptide that specifically uncouples pParkin dephosphorylation from pUb dephosphorylation, one can sustain pParkin signaling to enhance mitophagy while safeguarding broader cellular Ub signaling and homeostasis.

The PINK1-Parkin signaling is deeply implicated in Parkinson’s disease and other neurodegenerative disorders (1, 46). PP2A dysregulation is a recognized feature of Parkinson’s disease pathology, notably influencing the phosphorylation status of α-synuclein and Tau (47, 48). Our findings expand the PP2A substrate repertoire to include two critical mitophagy-related proteins. Currently, pharmacological strategies to enhance neuroprotective mitophagy primarily focus on inhibiting the deubiquitinase USP30 or employing small-molecule allosteric activators of Parkin (49, 50). Complementary to these approaches, the targeted disruption of the PP2A-Parkin interface using rationally designed peptides offers a novel, highly specific therapeutic avenue.

This approach selectively enhances neuroprotective mitophagy in the context of neurodegenerative disease, bypassing the pleiotropic side effects associated with broad-spectrum phosphatase inhibitors.

Despite the mechanistic insights, our study has several limitations that warrant future investigation. First, while AlphaFold modeling and functional validations support the non-canonical interface, an experimentally determined high-resolution structure of the PP2A–B55α– Parkin complex will be essential to resolve the precise atomic details of their interactions.

Second, the potential impacts of B55α manipulation and the competitive peptide on downstream TBK1–OPTN signaling and receptor-mediated (e.g., NIX/BNIP3) mitophagy have not been profiled. Directly assessing these interconnected pathways will be important for delineating the broader cellular consequences and establishing the precise degree of pathway selectivity. Third, although our rationally designed peptide demonstrates clear specificity and efficacy in cellular assays, its binding affinity could be enhanced, and its *in vivo* pharmacokinetic profile remains to be assessed. Advancing this proof-of-concept peptide into a viable therapeutic lead will require pharmacological optimization, such as macrocyclization, backbone modification, and the targeted incorporation of unnatural amino acids.

## Materials and Methods

### Recombinant proteins and *in vitro* assays

Human ubiquitin, Parkin constructs (full-length and Ubl), PINK1, PP1, and PP2A subunits (Aα, B55α, and Cα) were expressed and purified as detailed in *SI Appendix*. *In vitro* dephosphorylation assays were analyzed using Phos-tag SDS-PAGE, intact protein LC-MS, and ^1^H-^15^N HSQC / ^19^F NMR spectroscopy. Identification of the PP2A regulatory subunit from HeLa cell lysates was conducted via BS^3^ cross-linking followed by nanoLC-MS/MS on an Orbitrap mass spectrometer. Synthetic FITC-labeled Parkin peptides (PP#1–4) and the disulfide-constrained B55α β-hairpin peptide were synthesized (>95% purity) and assayed for direct binding using fluorescence polarization (FP).

### Cellular assays, microscopy, and immunoblotting

HeLa, YFP-Parkin HeLa, and SH-SY5Y cells were transfected with plasmids or siRNAs and subjected to mitochondrial depolarization (CCCP or oligomycin/antimycin A) in the presence or absence of competitive peptides. Mitophagic flux and mitochondrial translocation were assessed by quantitative immunofluorescence confocal microscopy (TIM23, TOM20, and Mito-QC reporter), JC-1 membrane potential assays, and quantitative Western blotting.

### Structural modeling, md simulations, and data availability

Complex modeling of PP2A–B55α bound to phosphorylated Parkin Ubl (residues 1–88) containing active-site Μn^2+^ cofactors was carried out using AlphaFold 3 (36), followed by 500-ns all-atom molecular dynamics simulations using AMBER24 (51). The structural model is deposited in Zenodo (DOI: 10.5281/zenodo.23035474). Statistical comparisons were performed with GraphPad Prism using ANOVA or Student’s *t*-test. Comprehensive experimental protocols, antibody dilutions, siRNA sequences, and analytical parameters are described in *SI Appendix*, *Materials and Methods*.

## Supporting information

Detailed Materials and Methods; Figures S1 to S8; Tables S1

## Acknowledgments

The work has been supported by Fundamental and Interdisciplinary Disciplines Breakthrough Plan of the Ministry of Education of China (JYB2025XDXM502), National Key R&D Program of China (2023YFF1204400), and Zhejiang Provincial Natural Science Foundation of China (LZ26H310003). We acknowledge the NMR facility at the National Center for Protein Sciences at Peking University for their technical support and assistance with data collection. We thank Dr. Sanhua Fang in the Core Facilities of Zhejiang University School of Medicine for technical assistance, and Prof. Luhua Lai for the use of BioTek instrument and Mr. Shuchen Zhong for assistance in FP measurement. The schematic graph was created with BioRender.com.

## References

1. J. Li et al., PINK1/Parkin-mediated mitophagy in neurodegenerative diseases. Ageing Res Rev 84, 101817 (2023).

2. D. P. Narendra, R. J. Youle, The role of PINK1-Parkin in mitochondrial quality control. Nat Cell Biol 26, 1639–1651 (2024).

3. S. M. Jin et al., Mitochondrial membrane potential regulates PINK1 import and proteolytic destabilization by PARL. J Cell Biol 191, 933–942 (2010).

4. L. A. Kane et al., PINK1 phosphorylates ubiquitin to activate Parkin E3 ubiquitin ligase activity. J Cell Biol 205, 143–153 (2014).

5. M. Lazarou et al., The ubiquitin kinase PINK1 recruits autophagy receptors to induce mitophagy. Nature 524, 309–314 (2015).

6. A. Kazlauskaite et al., Parkin is activated by PINK1-dependent phosphorylation of ubiquitin at Ser65. Biochem J 460, 127–139 (2014).

7. V. Sauvé et al., A Ubl/ubiquitin switch in the activation of Parkin. Embo j 34, 2492–2505 (2015).

8. T. Wauer, M. Simicek, A. Schubert, D. Komander, Mechanism of phospho-ubiquitin-induced PARKIN activation. Nature 524, 370–374 (2015).

9. C. Gladkova, S. L. Maslen, J. M. Skehel, D. Komander, Mechanism of parkin activation by PINK1. Nature 559, 410–414 (2018).

10. D. Narendra, A. Tanaka, D. F. Suen, R. J. Youle, Parkin is recruited selectively to impaired mitochondria and promotes their autophagy. J Cell Biol 183, 795–803 (2008).

11. J. Chakraborty et al., USP14 inhibition corrects an in vivo model of impaired mitophagy. EMBO Mol Med 10 (2018).

12. T. Cornelissen et al., The deubiquitinase USP15 antagonizes Parkin-mediated mitochondrial ubiquitination and mitophagy. Hum Mol Genet 23, 5227–5242 (2014).

13. J. R. Liang et al., USP30 deubiquitylates mitochondrial Parkin substrates and restricts apoptotic cell death. EMBO Rep 16, 618–627 (2015).

14. C. E. Wall et al., PPEF2 Opposes PINK1-Mediated Mitochondrial Quality Control by Dephosphorylating Ubiquitin. Cell Rep 29, 3280–3292.e3287 (2019).

15. L. Wang et al., PTEN-L is a novel protein phosphatase for ubiquitin dephosphorylation to inhibit PINK1-Parkin-mediated mitophagy. Cell Res 28, 787–802 (2018).

16. C. Chen et al., Elevated ubiquitin phosphorylation by PINK1 contributes to proteasomal impairment and promotes neurodegeneration. Elife 14 (2025).

17. C. Tang, W. P. Zhang, How Phosphorylation by PINK1 Remodels the Ubiquitin System: A Perspective from Structure and Dynamics. Biochemistry 59, 26–33 (2020).

18. S. Torres-Odio et al., Progression of pathology in PINK1-deficient mouse brain from splicing via ubiquitination, ER stress, and mitophagy changes to neuroinflammation. J Neuroinflammation 14, 154 (2017).

19. X. Dong et al., Ubiquitin S65 phosphorylation engenders a pH-sensitive conformational switch. Proc Natl Acad Sci U S A 114, 6770–6775 (2017).

20. S. S. Shekhawat, G. H. Pham, J. Prabakaran, E. R. Strieter, Simultaneous detection of distinct ubiquitin chain topologies by 19F NMR. ACS Chem Biol 9, 2229–2236 (2014).

21. S. X. Ye et al., Ubiquitin is double-phosphorylated by PINK1 for enhanced pH-sensitivity of conformational switch. Protein Cell 10, 908–913 (2019).

22. W. Li et al., Cantharidin, a potent and selective PP2A inhibitor, induces an oxidative stress-independent growth inhibition of pancreatic cancer cells through G2/M cell-cycle arrest and apoptosis. Cancer Sci 101, 1226–1233 (2010).

23. C. Bialojan, A. Takai, Inhibitory effect of a marine-sponge toxin, okadaic acid, on protein phosphatases. Specificity and kinetics. Biochem J 256, 283–290 (1988).

24. C. Gladkova et al., An invisible ubiquitin conformation is required for efficient phosphorylation by PINK1. Embo j 36, 3555–3572 (2017).

25. U. S. Cho, W. Xu, Crystal structure of a protein phosphatase 2A heterotrimeric holoenzyme. Nature 445, 53–57 (2007).

26. N. Wlodarchak, Y. Xing, PP2A as a master regulator of the cell cycle. Crit Rev Biochem Mol Biol 51, 162–184 (2016).

27. Y. Shi, Assembly and structure of protein phosphatase 2A. Sci China C Life Sci 52, 135– 146 (2009).

28. T. Kruse et al., Mechanisms of site-specific dephosphorylation and kinase opposition imposed by PP2A regulatory subunits. Embo j 39, e103695 (2020).

29. P. Amin et al., PP2A-B55: substrates and regulators in the control of cellular functions. Oncogene 41, 1–14 (2022).

30. Y. Xu, Y. Chen, P. Zhang, P. D. Jeffrey, Y. Shi, Structure of a protein phosphatase 2A holoenzyme: insights into B55-mediated Tau dephosphorylation. Mol Cell 31, 873–885 (2008).

31. F. Wachter, R. P. Nowak, S. Ficarro, J. Marto, E. S. Fischer, Structural characterization of methylation-independent PP2A assembly guides alphafold2Multimer prediction of family-wide PP2A complexes. J Biol Chem 300, 107268 (2024).

32. C. Kondapalli et al., PINK1 is activated by mitochondrial membrane potential depolarization and stimulates Parkin E3 ligase activity by phosphorylating Serine 65. Open Biol 2, 120080 (2012).

33. A. D. Baudot, M. Haller, M. Mrschtik, S. W. Tait, K. M. Ryan, Using enhanced-mitophagy to measure autophagic flux. Methods 75, 105–111 (2015).

34. S. A. Sarraf et al., Landscape of the PARKIN-dependent ubiquitylome in response to mitochondrial depolarization. Nature 496, 372–376 (2013).

35. T. G. McWilliams et al., mito-QC illuminates mitophagy and mitochondrial architecture in vivo. J Cell Biol 214, 333–345 (2016).

36. J. Abramson et al., Accurate structure prediction of biomolecular interactions with AlphaFold 3. Nature 630, 493–500 (2024).

37. H. Fowle et al., PP2A/B55α substrate recruitment as defined by the retinoblastoma-related protein p107. Elife 10 (2021).

38. S. K. R. Padi, R. J. Godek, W. Peti, R. Page, Cryo-EM structures of PP2A:B55 with p107 and Eya3 define substrate recruitment. Nat Struct Mol Biol 32, 1373–1382 (2025).

39. S. Shi et al., Cryo-EM structures reveal the PP2A-B55α and Eya3 interaction that can be disrupted by a peptide inhibitor. J Biol Chem 301, 110287 (2025).

40. V. Cianfanelli et al., The PP2A-B55α phosphatase is a master regulator of mitochondrial degradation and biogenesis. Sci Adv 11, eadw7376 (2025).

41. R. Cao et al., Molecular mechanism of PP2A/B55α phosphatase inhibition by IER5. Cell Chem Biol 32, 631–642.e637 (2025).

42. J. M. Heo, A. Ordureau, J. A. Paulo, J. Rinehart, J. W. Harper, The PINK1-PARKIN Mitochondrial Ubiquitylation Pathway Drives a Program of OPTN/NDP52 Recruitment and TBK1 Activation to Promote Mitophagy. Mol Cell 60, 7–20 (2015).

43. I. Novak et al., Nix is a selective autophagy receptor for mitochondrial clearance. EMBO Rep 11, 45–51 (2010).

44. L. P. Poole, A. Bock-Hughes, D. E. Berardi, K. F. Macleod, ULK1 promotes mitophagy via phosphorylation and stabilization of BNIP3. Sci Rep 11, 20526 (2021).

45. F. C. Fiesel et al., (Patho-)physiological relevance of PINK1-dependent ubiquitin phosphorylation. EMBO Rep 16, 1114–1130 (2015).

46. M. S. D’Arcy, Mitophagy in health and disease. Molecular mechanisms, regulatory pathways, and therapeutic implications. Apoptosis 29, 1415–1428 (2024).

47. K. W. Lee et al., Enhanced phosphatase activity attenuates α-synucleinopathy in a mouse model. J Neurosci 31, 6963–6971 (2011).

48. P. Javadpour, L. Dargahi, A. Ahmadiani, R. Ghasemi, To be or not to be: PP2A as a dual player in CNS functions, its role in neurodegeneration, and its interaction with brain insulin signaling. Cell Mol Life Sci 76, 2277–2297 (2019).

49. V. Sauvé et al., Activation of parkin by a molecular glue. Nat Commun 15, 7707 (2024).

50. E. V. Rusilowicz-Jones et al., Benchmarking a highly selective USP30 inhibitor for enhancement of mitophagy and pexophagy. Life Sci Alliance 5 (2022).

51. R. Salomon-Ferrer, A. W. Götz, D. Poole, S. Le Grand, R. C. Walker, Routine Microsecond Molecular Dynamics Simulations with AMBER on GPUs. 2. Explicit Solvent Particle Mesh Ewald. J Chem Theory Comput 9, 3878–3888 (2013).

