## Supplementary material for "PP2A-B55α Terminates Mitophagy by Selectively Dephosphorylating Parkin via a Non-Canonical Interface": Detailed Materials and Methods; Figures S1 to S8; Tables S1

**This PDF file includes:**

Detailed Materials and Methods
Figures S1 to S8
Tables S1

### Materials and Methods

#### Protein sample preparation

K48C, T66V/L67N, and N-terminal FLAG mutations were introduced into the human ubiquitin gene using the QuikChange method, and the resulting genes were cloned into a pET11a vector. Except for FLAG-Ub, the expression and purification of the ubiquitin proteins followed the established protocol (1). As for FLAG-Ub, the protein was purified with Source-Q (GE Healthcare) and Sephacryl S100 columns (GE Healthcare) in tandem. For uniform $^{15}$N-labeling, BL21-star cells transformed with the ubiquitin plasmid were grown in M9-minimal medium with $^{15}NH_4Cl$ (Cambridge Isotope Laboratory) as the sole nitrogen source.

PP1α was cloned into a pET-11a vector with an 8×His tag appended at its N-terminus. BL21-star cells were used for protein expression, and were grown in LB medium at 37 ºC and induced by 0.2 mM IPTG for 20 hours at 18 ºC before harvesting. The protein was purified with Ni-NTA Superflow affinity column (GE Healthcare) and Sephacryl S100 columns (GE Healthcare) in tandem. PP2A scaffolding subunit Aα and regulatory subunit B55α were cloned into the pGEX-6P-1vector, respectively. Cells were grown in LB medium at 37 °C and induced by 0.2 mM IPTG for 20 hours at 18 °C before harvesting. The proteins were purified using Glutathione Sepharose 4B (GE Healthcare) and Sephacryl S100 columns (GE Healthcare) in tandem.

PP2A catalytic subunit Cα was cloned into the pFastBac1 vector with a 10×His tag appended at the N terminus and was expressed in Sf9 cells (Gibco, catalog number 12659017). The cells were cultured at 27 ℃ for 60 h after being infected with baculoviruses using the Bac-to-Bac system (Life Technologies). The recombinant proteins were purified using a Ni-NTA Superflow affinity column (GE Healthcare) and Source-Q (GE Healthcare) in tandem.

Parkin was cloned into a pET11a vector, with a His-SUMO tag appended at the N-terminus. BL21-star cells were used to express proteins, and were grown in LB medium at 37 °C and induced by 0.1 mM IPTG and 200 mM $ZnCl_2$ for 20 hours at 18 °C before harvesting. The protein was purified with the Ni-NTA Superflow affinity column (GE Healthcare) and Sephacryl S100 column (GE Healthcare) in tandem. The isolated Parkin Ubl domain (residues 1-76 from the full-length human Parkin) was cloned into a pGEX-6P-1 vector. The protein was purified with a Glutathione Sepharose 4B column (GE Healthcare) and Sephacryl S100 column (GE Healthcare) in tandem. The His-Sumo and GST tags at the N-terminus of the desired proteins were removed by incubation with PreScission Protease (Beyotime Biotechnology) at room temperature for 3 h.

PINK1 from body louse was prepared as previously described (1). To phosphorylate ubiquitin and Parkin, PINK1, ubiquitin (or Parkin), $MgCl_2$, and ATP were mixed at a molar ratio of 1:100:5,000:5,000 in 20 mM Tris•HCl buffer (pH 8.0, containing 1 mM DTT). The reaction proceeded overnight at room temperature. To phosphorylate the isolated Parkin Ubl domain, the reaction was carried out with the same molar ratio and conditions, but only for 30 min. The resulting proteins, pUb, pParkin, and pUbl, were purified on a Source-Q column (GE Healthcare), buffer-exchanged; the modifications were confirmed with a Vion IMS QTOF Mass Spectrometer (Waters).

3-Bromo-1,1,1-trifluoroacetone (BFA) (J&K Scientific, catalog number 312226) was dissolved in DMSO to 5 M concentration as a stock solution. The K48C mutant of ubiquitin was desalted (HiPrep 26/10 Desalting column, GE Healthcare), and mixed with BFA at a molar ratio of 1:10 in 20 mM pH 7.4 HEPES buffer. The conjugation reaction was performed at room temperature for 2 h. The product was further purified using the desalting column, and the modification was confirmed with a Vion IMS QTOF Mass Spectrometer (Waters).

**Phos-tag gel**

To evaluate the phosphorylation level of the full-length Parkin, the protein samples were resolved on a 12% polyacrylamide gel containing 50 μM Phos-tag acrylamide (Wako Chemicals, catalog number AAL-107) and 100 μM $MnCl_2$. Tricine protein gel with 12% polyacrylamide containing 100 μM Phos-tag acrylamide and 200 μM $MnCl_2$ was used to assess the phosphorylation level of the isolated Parkin Ubl domain and Ub. For sample preparation, 100 μM pParkin, pUb, or pUbl was incubated in enzyme buffer (50 mM Tris, pH 7.4, 150 mM NaCl, 1mM $MnCl_2$) with 0.5 μM PP2A (core enzyme or holoenzyme, as noted in the Results) or buffer only at the room temperature, with the incubation time noted in the Results.

**Mass spectrometry identification of PP2A regulatory subunit**

To identify the PP2A regulatory subunit, an excess amount of HeLa cell lysate was prepared by manual grinding in lysis buffer (50 mM HEPES, pH 7.4, 150 mM NaCl) in the presence of protease inhibitors (Coolaber, catalog number SL1086). The lysate was mixed with recombinantly purified FLAG-pUb and GST-PP2A Aα subunit. Subsequently, 4 mM freshly prepared $BS_3$ (Thermo Fisher, catalog number 21580) was added. The cross-linking reaction was conducted at room temperature with gentle mixing and was quenched after 45 min by adding Tris-HCl buffer (pH 8.0) to a final concentration of 20 mM. The proteins were pooled onto GST affinity resin (GE Healthcare), incubated for 2 h at 4 °C, washed three times with lysis buffer, and eluted with lysis buffer containing 20 mM L-Glutathione reduced (Aladdin, catalog number G105426). Eluted proteins were incubated with anti-Flag magnetic beads (Bimake, catalog number B26102) for 2 h at 4 °C. Subsequently, the beads were washed three times in lysis buffer and eluted by the addition of 0.3 mg/mL poly-FLAG peptide (Bimake, catalog number B23111). The eluted proteins were subjected to trypsin digestion.

The resulting peptide mixture was subjected to an Easy-nLC 1200 nano HPLC (Thermo Scientific) coupled to a Q Exactive Orbitrap mass spectrometer (Thermo Scientific). Samples were resolved on an in-house packed C18 RP-trap column (150 μm i.d. × 3 cm) and a C18 capillary column (150 μm i.d. × 15 cm) with ReproSil-Pur C18-AQ particles (1.9 μm, 120 Å). Mobile phase A contained 0.1% FA in HPLC $H_2O$, and mobile phase B contained 80% acetonitrile, 0.1% FA, and 20% HPLC $H_2O$. The flow rate was 300 nL/min, and the elution gradient was as follows: 5-10% buffer B for 1 min, 10-40% buffer B for 95 min, 40-60% buffer B for 8 min, 60-100% buffer B for 1 min, and finally 100% buffer B for 15 min. The mass spectrometer was operated in data-dependent mode with one full MS scan at $R$ = 70,000 (m/z = 200), followed by HCD MS/MS scans at $R$ = 17,500 (m/z = 200), NCE = 30, with an isolation width of 1.8 m/z. The AGC targets for the $MS_1$ and $MS_2$ scans were 3e+6 and 1e+5, respectively; the maximum injection times for $MS_1$ and $MS_2$ were 50 ms and 100 ms, respectively. The pFind Studio 3 software (2) was used to identify the interacting peptides. Search parameters were set as follows: Cys carbamidomethyl as a static modification and Met oxidation as a variable modification, precursor mass tolerance of 20 ppm, fragment mass tolerance of 20 ppm, full tryptic cleavage constraints allowing up to 3 missed cleavages, and a spectral false identification rate ≤1%.

**NMR spectroscopy analysis**

For $^{1}H$-$^{15}N$ HSQC NMR experiments, samples were prepared in 20 mM HEPES buffer (150 mM NaCl, pH 7.4, with 5% $D_2O$), and NMR data were acquired at 298 K on a Bruker 600 MHz spectrometer equipped with a Z-gradient cryogenic probe. The data were processed using NMRPipe (3) and analyzed using

NMRViewJ (4). The protein samples used in $^{19}$F NMR were prepared in 50 mM Tris•HCl buffer (150 mM NaCl, 1 mM $MnCl_2$, and pH 7.4, with 5% $D_2O$). NMR experiments were recorded at 298 K on a 600 MHz spectrometer (Bruker) equipped with a triple-resonance cryogenic probe. The $^{19}$F base frequency is 564.39 MHz, with the 0 ppm set for the peak from trichlorofluoromethane. The peak volumes in the one-dimensional $^{19}$F NMR spectra were obtained with Bruker TopSpin Version 3.5 and NMRViewJ (4). Since the $^{19}$F peaks of the phosphorylated and unphosphorylated Ub T66V/L67N mutant could not be completely distinguished, the relative populations of the three peaks were fitted using the Lorentz function with OriginPro 2019 (OriginLab).

For the dephosphorylation reaction, HeLa cells were harvested, washed twice with ice-cold 1x PBS buffer, and lysed at 4℃ in the RIPA buffer (Beyotime Biotechnology, catalog number P0013C) with protease inhibitors (Coolaber, catalog number SL1086). Total protein concentrations were determined using a BCA protein assay kit (Beyotime Biotechnology, catalog number P0012S). HeLa cell lysate was added to 100 µM $^{15}$N-labeled pUb at a 1:1 (w/w) ratio. The reaction proceeded at room temperature for 4 h. HeLa cell lysate was also added to 100 µM $^{19}$F-labeled pUb at a 1:1 (w/w) ratio for the dephosphorylation reaction at room temperature. To inhibit the dephosphorylation reaction, 1 µM okadaic acid (Aladdin, catalog number O275664) or 10 µM Cantharidin (Aladdin, catalog number C111020) was added to 100 µM $^{19}$F-labeled pUb, before mixing with the cell lysate at a 1:1 (w/w) ratio. A series of 1D $^{19}$F NMR data were collected at room temperature to monitor the dephosphorylation reaction in real-time. In addition, recombinantly purified 0.5 µM PP2A (core enzyme or holoenzyme) or 0.5 µM PP1 core enzyme was added into 100 µM $^{19}$F-labeled pUb for dephosphorylation.

**Cell lines and treatment of the cells**

HeLa cells stably expressing YFP-Parkin were kind gifts from Richard Youle as previously described (5). The SH-SY5Y cells (CAS Center for Excellence in Molecular Cell Science, China) were seeded onto poly-D-lysine (Sigma-Aldrich, catalog number P7405) coated glass-bottom dishes (Cellvis, catalog number D29-20-1-N) for live-cell imaging. The SH-SY5Y cells, regular HeLa cells (CAS Center for Excellence in Molecular Cell Science, China), and YFP-Parkin HeLa cells were grown in Dulbecco's modified essential medium (DMEM, Gibco, catalog number 11965092) supplemented with 10% heat-inactivated fetal bovine serum (Gibco, catalog number 2275120) at 37 °C, in the atmosphere of 5% $CO_2$.

**Cell transfection and treatment**

Transient transfections of plasmids were performed using Lipofectamine 2000 (Thermo Fisher Scientific, catalog number 11668019) or Lipofectamine 3000 (Thermo Fisher Scientific, catalog number L3000015). For plasmid construction, Parkin and PP2A-Cα-IRES-Aα were cloned into a pcDNA3.1 vector (PPP1CA/PP1α). PP2A-Cα alone was cloned into a pcDNA3.1-mCherry vector. A 3×Flag-tagged PP1 expression construct was generated using the pcDNA3.1 vector. Untagged full-length Parkin mutants R72A, R75A, and R72A/R75A were generated from the wild-type pcDNA3.1-Parkin construct using QuikChange mutagenesis. B55α constructs (wild-type and mutants) were cloned into a pcDNA3.1 vector with a C-terminal 3×Flag tag. The Mito-QC plasmid (6) was constructed by cloning EGFP and the mitochondrial targeting sequence of human Fis1 (residues 101–152) into a pmCherry-C1 vector.

When HeLa or YFP-Parkin HeLa cells reached ~90% confluency, plasmids including pcDNA3.1-PP2A-Cα-IRES-Aα, pcDNA3.1-PP1α, pcDNA3.1-Parkin (wild-type, Parkin-R72A, Parkin-R75A, and Parkin-R72A/R75A), pcDNA3.1-PP2A-Cα-mCherry, or the B55α-3×Flag constructs (wild-type, B55α-E91A, B55α-E93A, B55α-E94A, B55α-3A, and B55α-F84A generated via QuikChange mutagenesis) were transfected. Empty vectors were used as controls to ensure equal amounts of total DNA (typically 2-4 µg per well in 6-

well plates). For live-cell mitophagy assays, SH-SY5Y cells were transfected with the Mito-QC plasmid using Lipofectamine 3000.

For gene knockdown, siRNA transfections were achieved using Lipofectamine RNAiMAX (Thermo Fisher Scientific, catalog number 13778075) at a final concentration of 50 nM. The siRNA sequences were as follows: PP2A Cα siRNA (5'-GGAACUUGACGAUACUCUA-3'), B55α siRNA #1 (siB55α#1; 5′-GCAGAUGAUUUGCGGAUUAAU-3′, knockdown data presented in Fig. S3F), B55α siRNA #2 (siB55α#2; 5′-GCCGUAGCUACUACAAACATT-3′, knockdown data presented in Fig. 3D), PR130 siRNA (5′-GCGCCAUUGUCCAUAAACATT-3′), and control siRNA (5'-UUCUCCGAACGUGUCACGU-3'). For co-transfection rescue experiments, cells at ~60% confluency were transfected with specific siRNAs alongside the indicated plasmids.

To induce mitophagy, cells were cultured for 24 h post-plasmid transfection or 48 h post-siRNA transfection, followed by treatment with 10 μM CCCP (Sigma-Aldrich, catalog number C2759) or 250 nM oligomycin (Abcam, catalog number ab141829) combined with 250 nM antimycin A (Sigma-Aldrich, catalog number A8674) (O/A).

**Peptide synthesis and treatment**

A series of short peptides derived from the C-terminus of the Parkin Ubl domain N-terminally conjugated with FITC were commercially synthesized by TGpeptide Biotechnology. The peptide panel included the wild-type mimetic PP#1 (residues 71-77, sequence: QRPWRKG) and three variants with single amino acid substitutions at the Trp74 position: PP#2 (W74Y), PP#3 (W74L), and PP#4 (W74G). Additionally, a peptide corresponding to the protruding β-hairpin of the blade 4 in B55α (residues 79–97, sequence: SHEPEFDYLKSLEIEEKIN) was synthesized. To promote conformational stability and constrain the native β-hairpin fold in solution, a pair of cysteine substitutions (H80C and I96C) was introduced to allow the formation of an intramolecular disulfide bridge. All peptides were purified to >95% purity via HPLC.

For intracellular peptide delivery, the FITC-conjugated peptides were directly added to the cell culture medium at a final concentration of 50 μM. After 3 h of incubation at 37 °C, the cells were treated with 10 μM CCCP to induce mitophagy. The intracellular distribution of the peptides and their effects on pParkin preservation and mitophagic flux were subsequently evaluated by Western blotting and immunofluorescence.

**Western blotting analysis**

Cells were washed 3 times with ice-cold 1×PBS buffer and lysed with the RIPA buffer with protease (Coolaber, catalog number SL1086) and phosphatase inhibitors (Coolaber, catalog number SL1087) present. The cell lysate was quantified using the BCA protein assay kit and was reduced and boiled before loading onto a 12% polyacrylamide gel. The proteins were transferred from the gel to PVDF membranes. The membranes were blocked with 5% nonfat milk in TBS buffer containing 0.05% Tween-20 (TBST) for 1 h, incubated with primary antibodies in the blocking buffer (TBST buffer with 5% nonfat milk) overnight at 4ºC, and then HRP-conjugated secondary antibodies (diluted 1:2000 in blocking buffer) for 2 h at room temperature. Protein quantity was extrapolated based on HRP activity using the Clarity™ Western ECL Substrate (Bio-Rad Laboratories, catalog number 1705060), measured on a ChemiDoc XRS+ (BioRad), and analyzed using the Image Lab software (Bio-Rad Laboratories).

The primary antibodies used for Western blotting included those for Parkin (Cell Signaling Technology, catalog number 2132S, diluted at 1:1000 ratio), phospho-Parkin (Ser65) (Cell Signaling Technology, catalog number 36866S, diluted at 1:500 ratio), ubiquitin (Beyotime Biotechnology, catalog number AF1705, diluted at 1:1000 ratio), phospho-ubiquitin (Ser65) (Merck Millipore, catalog number ABS1513-I, diluted at 1:500 ratio), PP2A Cα (Invitrogen, catalog number MA5-32920, diluted at 1:2000 ratio), PP2A B55α (PPP2R2A, Affinity Biosciences, catalog number DF6908, diluted at 1:1000 ratio), TIM23 (Proteintech Group, catalog number 11123-1-AP, diluted at 1:5000 ratio), PP1α (Proteintech, catalog number 85445-3-RR, diluted at 1:5000 ratio), PR130 (PPP2R3A, Genetex, catalog number GTX114655, diluted at 1:1000 ratio), GAPDH (Proteintech, catalog number 60004-1-Ig, diluted at 1:20000 ratio). Secondary antibodies were purchased from Beyotime Biotechnology, including HRP-conjugated goat anti-rabbit IgG (catalog number A0208) and HRP-conjugated goat anti-mouse IgG (catalog number A0216).

**Immunofluorescence imaging**

For fixed-cell immunofluorescence, following CCCP or O/A treatment, cells were washed with 1×PBS and fixed with 4% paraformaldehyde in PBS for 15 min at 37 °C. After three washes with 1×PBS, cells were permeabilized with 0.1% Triton X-100 in PBS for 20 min. Cells were then washed and blocked with 5% normal donkey serum (Yeasen Biotechnology, catalog number 36116ES03) in PBS for 30 min at room temperature. Primary antibodies were diluted in the blocking solution (1:200) and incubated overnight at 4 °C in a humidified chamber. Following three washes with PBS, cells were incubated with fluorophore-conjugated secondary antibodies, including Alexa Fluor 647-conjugated goat anti-mouse IgG (Thermo Fisher Scientific, A-21235) or Alexa Fluor 594-conjugated goat anti-rabbit IgG (Thermo Fisher Scientific, A-11080) (diluted 1:200), for 2 h at room temperature in the dark. Cells were washed three times with PBS and mounted using DAPI Fluoromount-G™ (Yeasen Biotechnology, catalog number 36308ES11). Images were acquired using a Laser Scanning Confocal Microscope (Nikon, A1R-si) or a STELLARIS 8 confocal microscope (Leica) with a 60× oil immersion objective. Primary antibodies included anti-TOM20 (Santa Cruz, SC-17764), anti-Flag (Proteintech Group, 66008-4-Ig), anti-Parkin (CST, catalog number 4211), and anti-TIM23 (Proteintech Group, 11123-1-AP). The colocalized areas of YFP-Parkin or Parkin and TOM20 or TIM23 fluorescence were quantified using Nikon NIS-Elements AR imaging software.

For live-cell imaging of mitophagy, SH-SY5Y cells expressing the Mito-QC reporter were imaged at 37 °C following various treatments. Images were acquired using an LSM880 inverted confocal microscope (Zeiss) equipped with a 63× oil objective. The excitation wavelengths were set at 488 nm for GFP and 548 nm for mCherry, with corresponding emission collected at 519 nm and 562 nm, respectively. For quantitative analysis, red-only puncta in each cell were counted as mito-lysosomes, and the total mitochondrial area of the corresponding cells was measured using ImageJ software.

For experiments evaluating the efficacy of competitive peptides, cells were immunostained for the inner mitochondrial membrane marker TIM23. To quantify mitochondrial clearance, the TIM23-positive mitochondrial area and the mean fluorescence intensity (MFI) per cell were measured using ImageJ/Fiji software (National Institutes of Health). A threshold was uniformly applied to all images to delineate the mitochondrial network, and the total area and MFI of the suprathreshold pixels were calculated.

**Mitochondrial membrane potential assay**

Mitochondrial membrane potential was assessed using a JC-1 mitochondrial membrane potential assay kit (Beyotime, catalog number C2006) in HeLa cells subjected to the indicated experimental conditions. Where indicated, cells were treated with 10 μM CCCP for 1 h before JC-1 staining. The JC-1 staining working solution was prepared according to the manufacturer's instructions and mixed with an equal volume of culture medium. Cells were incubated for 20 min at 37 °C. Following staining, cells were washed three times with PBS, and live-cell fluorescence images were acquired using a STELLARIS 8 confocal microscope (Leica). Images were analyzed using ImageJ software (NIH), and the ratio of red (aggregates) to green (monomer) JC-1 fluorescence intensity was calculated as an indicator of mitochondrial membrane potential.

**Fluorescence polarization (FP) binding assay**

Direct physical interaction between the B55α β-hairpin and the Parkin Ubl domain was evaluated by *in vitro* fluorescence polarization (FP) assays using the FITC-labeled peptide derived from the Parkin Ubl domain and the unlabeled synthetic B55α peptide. The FITC-labeled Parkin peptide was maintained at a concentration of 5 μM in the PBS buffer pH 7.4 and 0.01% (v/v) Tween-20 to minimize non-specific plate adsorption. Serial concentrations of the unlabeled B55α peptide were incubated in 96-well black, non-binding microplate for 10 min at room temperature protected from light. Fluorescence polarization signals were measured on a microplate reader (BioTek Synergy) with excitation at 485 nm and emission at 535 nm. Parallel and perpendicular fluorescence intensities were recorded, and millipolarization (mP) values were calculated. The titration was performed in triplicates, and binding curves were fitted to a standard one-site binding isotherm using Origin.

**Structural modeling and molecular dynamics simulations**

Structural models of the quaternary complex comprising the heterotrimeric PP2A holoenzyme and the phosphorylated Parkin Ubl domain were generated using AlphaFold3 (7). The modeled protein assembly included the catalytic subunit Cα (PP2Ac, Uniprot: P67775), the scaffolding subunit PR65A (residues 1–589, Uniprot: P30153), the regulatory subunit B55α (residues 1–447, Uniprot: P63151-1), the phosphorylated Parkin Ubl domain (Uniprot: O60260), and two $Mn^{2+}$ ions.

To determine the optimal substrate boundary, we systematically varied Parkin construct lengths, including the minimal Ubl domain (residues 1–78), extended linker (residues 1–99), and full-length Parkin (residues 1–465). Inclusion of full-length Parkin or extended disordered linker—where Ile142 in the RING0 domain is the first downstream structured residue—introduced excessive conformational entropy and steric noise during diffusion sampling, steering the substrate domain away from the catalytic cleft. Defining the substrate construct as Parkin residues 1–88 preserved the folded Ubl core (residues 1–71) and the functional basic C-terminal tail (residues 72–88) while eliminating entropic sampling penalties.

Phosphorylation at Ser65 and the presence of the active-site binuclear metal center were explicitly defined; control prediction runs omitting either the metal cofactors or the phosphoryl group failed to position the substrate near the active site. Structure generation was performed across five independent runs using distinct random seeds (yielding 5 models per run; 25 models in total). Models were evaluated based on the interface predicted TM-score (ipTM), inter-subunit Predicted Aligned Error (PAE) matrices between B55α and the Ubl domain, and catalytic geometry (defined by spatial proximity between the $pSer^{65}$ phosphoryl group and the binuclear catalytic metal center). Across all calculations, 15 out of 25 models (60%) converged on a consistent quaternary architecture. Atomic coordinates of the representative converged model are publicly accessible at Zendo with DOI of 10.5281/zenodo.23035474.

To prepare the coordinates, protonation states of titratable residues and phosphoryl group were adjusted to pH 7.4 with CHARMM-GUI, and neutralized with 0.15 M KCl (8). The protein was parameterized using the CHARMM36m force field (9). Hydrogen Mass Repartitioning (HMR) was applied to enable a 4-fs integration time step. Long-range electrostatic interactions were treated using the Particle Mesh Ewald (PME) method with a cutoff distance of 1.2 nm for both van der Waals and short-range electrostatic interactions with force-based switching at 1.0 nm. The temperature was maintained at 310 K using the

Langevin thermostat, and pressure was maintained at 1 bar using the Monte Carlo barostat. All-atom molecular dynamics (MD) simulations of PP2A-Parkin system were performed using the AMBER24 software package, with AMBER pmemd.cuda used to run the simulation for 500 ns (10). Post-simulation trajectories were aligned and analyzed using CPPTRAJ (11).

**Statistical analysis**

All data were processed using Prism 9.0 (GraphPad) and are presented as means ± SEM from at least three independent biological replicates ($n \geq 3$). For image-based quantifications, including the colocalization analysis, Mito-QC assays, and TIM23-positive area/mean fluorescence intensity measurements, 15-30 cells per condition were randomly selected and analyzed. Statistical significance between multi-group comparisons was determined by two-way analysis of variance (ANOVA) or one-way ANOVA. Comparisons between two independent groups were analyzed using an unpaired two-tailed Student's *t*-test. *P* values are indicated with asterisks with *$P < 0.05$, **$P < 0.01$, ***$P < 0.001$, ****$P < 0.0001$, and *N.S.* for not significant.


1. X. Dong *et al.*, Ubiquitin S65 phosphorylation engenders a pH-sensitive conformational switch. *Proc Natl Acad Sci U S A* **114**, 6770–6775 (2017).
2. H. Chi *et al.*, Comprehensive identification of peptides in tandem mass spectra using an efficient open search engine. *Nat Biotechnol* **36**, 1059–1061 (2018).
3. Delaglio F, Stephan Grzesiek, Geerten W. Vuister b, G. Z. r. J. P. d. a. A. Bax, NMRPipe: A multidimensional spectral processing system based on UNIX pipes. *Journal of Biomolecular NMR* **6**, 277–293 (1995).
4. Bruce A. Johnsona, R. A. Blevinsb, NMRView: A computer program for the visualization and analysis of NMR data. *Journal of Biomolecular NMR,* **4**, 603–614 (1994).
5. D. Narendra, A. Tanaka, D. F. Suen, R. J. Youle, Parkin is recruited selectively to impaired mitochondria and promotes their autophagy. *J Cell Biol* **183**, 795–803 (2008).
6. T. G. McWilliams *et al.*, mito-QC illuminates mitophagy and mitochondrial architecture in vivo. *J Cell Biol* **214**, 333–345 (2016).
7. J. Abramson *et al.*, Accurate structure prediction of biomolecular interactions with AlphaFold 3. *Nature* **630**, 493–500 (2024).
8. S. Jo, T. Kim, V. G. Iyer, W. Im, CHARMM-GUI: a web-based graphical user interface for CHARMM. *J Comput Chem* **29**, 1859–1865 (2008).
9. J. Huang *et al.*, CHARMM36m: an improved force field for folded and intrinsically disordered proteins. *Nat Methods* **14**, 71–73 (2017).
10. R. Salomon-Ferrer, A. W. Götz, D. Poole, S. Le Grand, R. C. Walker, Routine Microsecond Molecular Dynamics Simulations with AMBER on GPUs. 2. Explicit Solvent Particle Mesh Ewald. *J Chem Theory Comput* **9**, 3878–3888 (2013).
11. D. R. Roe, T. E. Cheatham, 3rd, PTRAJ and CPPTRAJ: Software for Processing and Analysis of Molecular Dynamics Trajectory Data. *J Chem Theory Comput* **9**, 3084–3095 (2013).

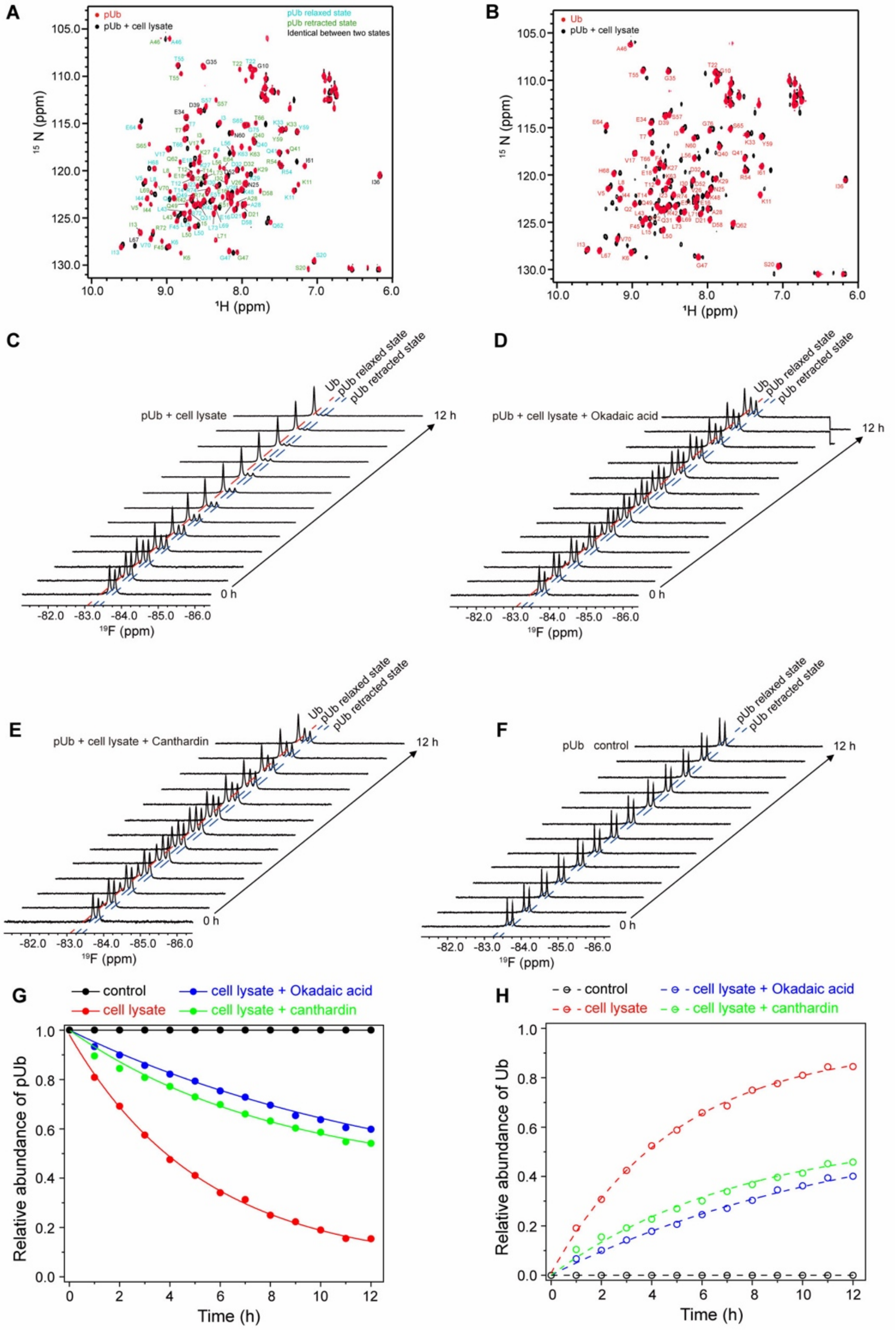


**Fig. S1 Identification of PP2A as a potential phosphatase for pUb using cell lysates and specific inhibitors. (A, B)** Overlay of $^{1}$H-$^{15}$N HSQC spectra of 100 μM $^{15}$N-labeled pUb treated with HeLa cell lysate (1:1 w/w total protein ratio) for 4 h. The treated spectrum is compared with untreated pUb **(A)** and unmodified ubiquitin **(B)**. Approximately 30-40% of pUb remained after lysate treatment. **(C-F)** Real-time $^{19}$F-NMR spectra of 100 μM $^{19}$F-labeled pUb (probe conjugated at K48C) treated with cell lysate **(C)**, lysate + 1 μM okadaic acid **(D)**, lysate + 10 μM cantharidin **(E)**, or buffer control **(F)** over 12 h. **(G, H)** Quantification of the relative abundance of pUb **(G)** and unmodified Ub **(H)** based on $^{19}$F peak intensities from **(C-F)**, normalized to time 0.

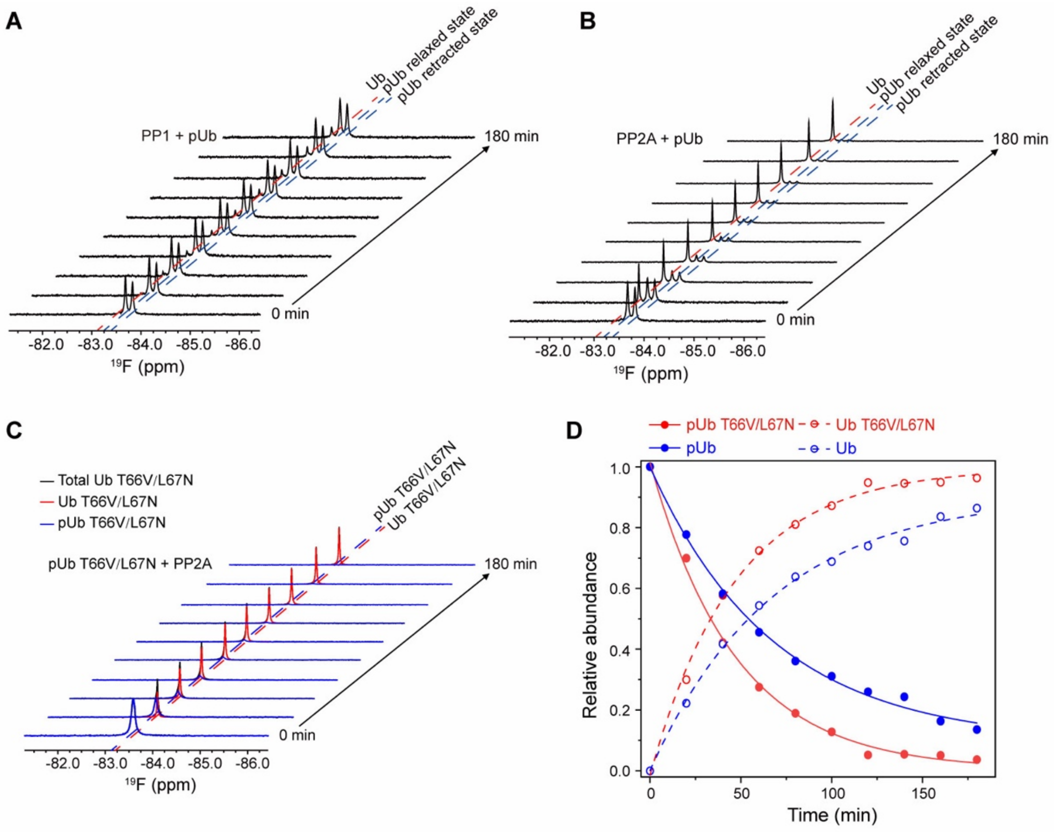


**Fig. S2. PP2A core enzyme preferentially dephosphorylates pUb in retracted state. (A, B)** Real-time $^{19}$F-NMR spectra of 100 μM pUb in the presence of 0.5 μM PP1 **(A)** or PP2A core enzyme **(B)**. **(C)** Real-time $^{19}$F-NMR spectra monitoring the dephosphorylation of 100 μM pUb T66V/L67N double mutant (fully adopting the retracted state) in the presence of 0.5 μM PP2A core enzyme. **(D)** Kinetic quantification of pUb and unmodified Ub relative abundance over time, extracted from the NMR spectra in (**B** and **C**). Data are normalized to the initial intensities at 0 min.

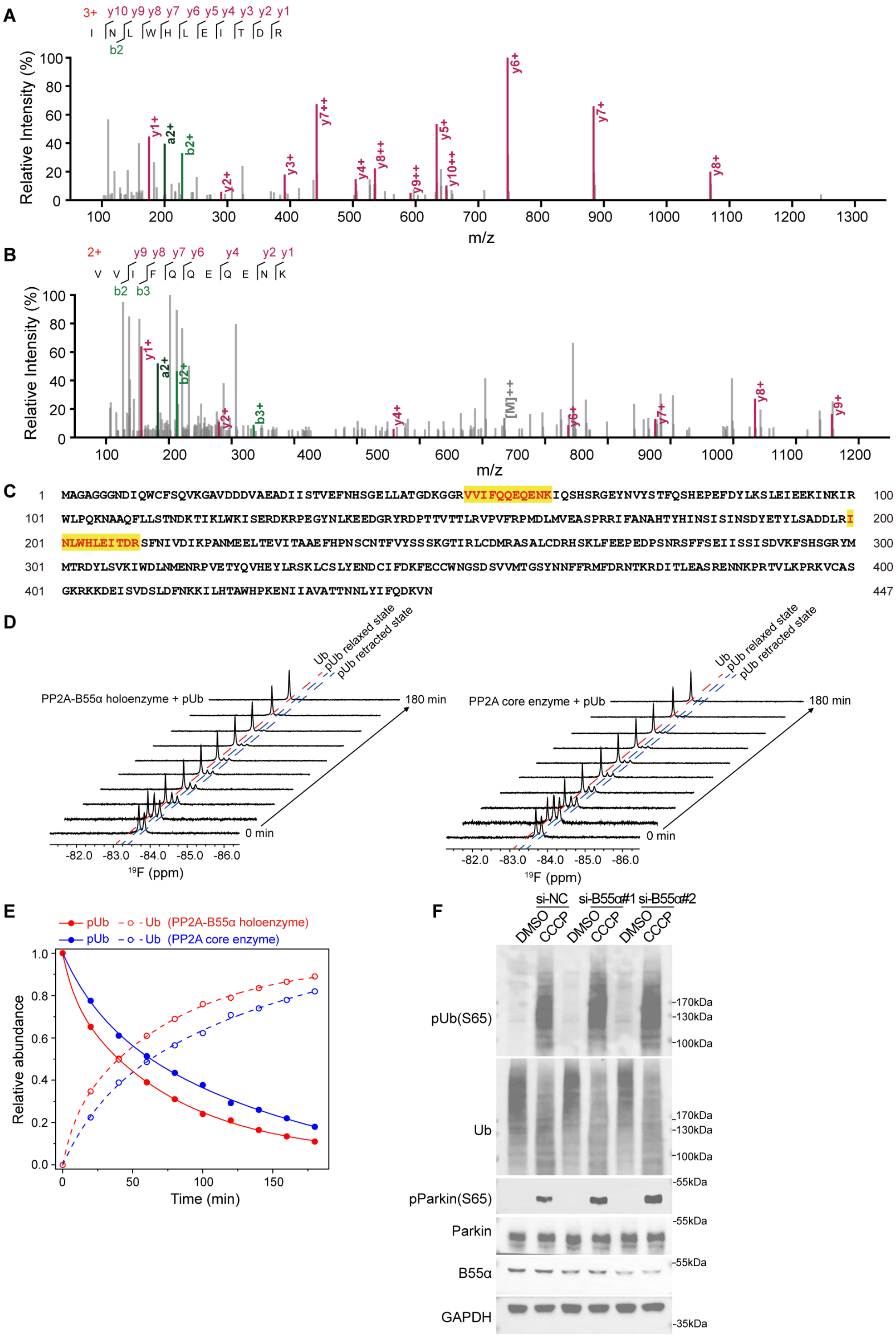


**Fig. S3. Identification and functional validation of B55α as the PP2A regulatory subunit. (A, B)** High-resolution LC-MS/MS spectra of representative peptides identified from the dual-affinity pulldown eluate. **(C)** Mapping of the identified peptides (highlighted in yellow and red) to the human B55α protein sequence. **(D)** Real-time $^{19}$F-NMR spectra monitoring the dephosphorylation of 100 μM pUb in the presence of the reconstituted PP2A-B55α holoenzyme (left) or the PP2A core enzyme (right). **(E)** Kinetic quantification of pUb and unmodified Ub relative abundance extracted from the NMR spectra in **(D)**. Data are normalized to the initial $^{19}F$ peak intensities at 0 min. The unmodified Ub peak intensity was calculated by subtracting the remaining pUb intensity from the total intensity. **(F)** Validation of the effect of B55α depletion using two independent siRNAs. HeLa cells were transfected with si-NC or either of two independent siRNAs targeting B55α (si-B55α#1 and si-B55α#2), followed by treatment with DMSO or CCCP (10 μM for 4 h). Cell lysates were analyzed by immunoblotting using the indicated antibodies, with GAPDH as a loading control.

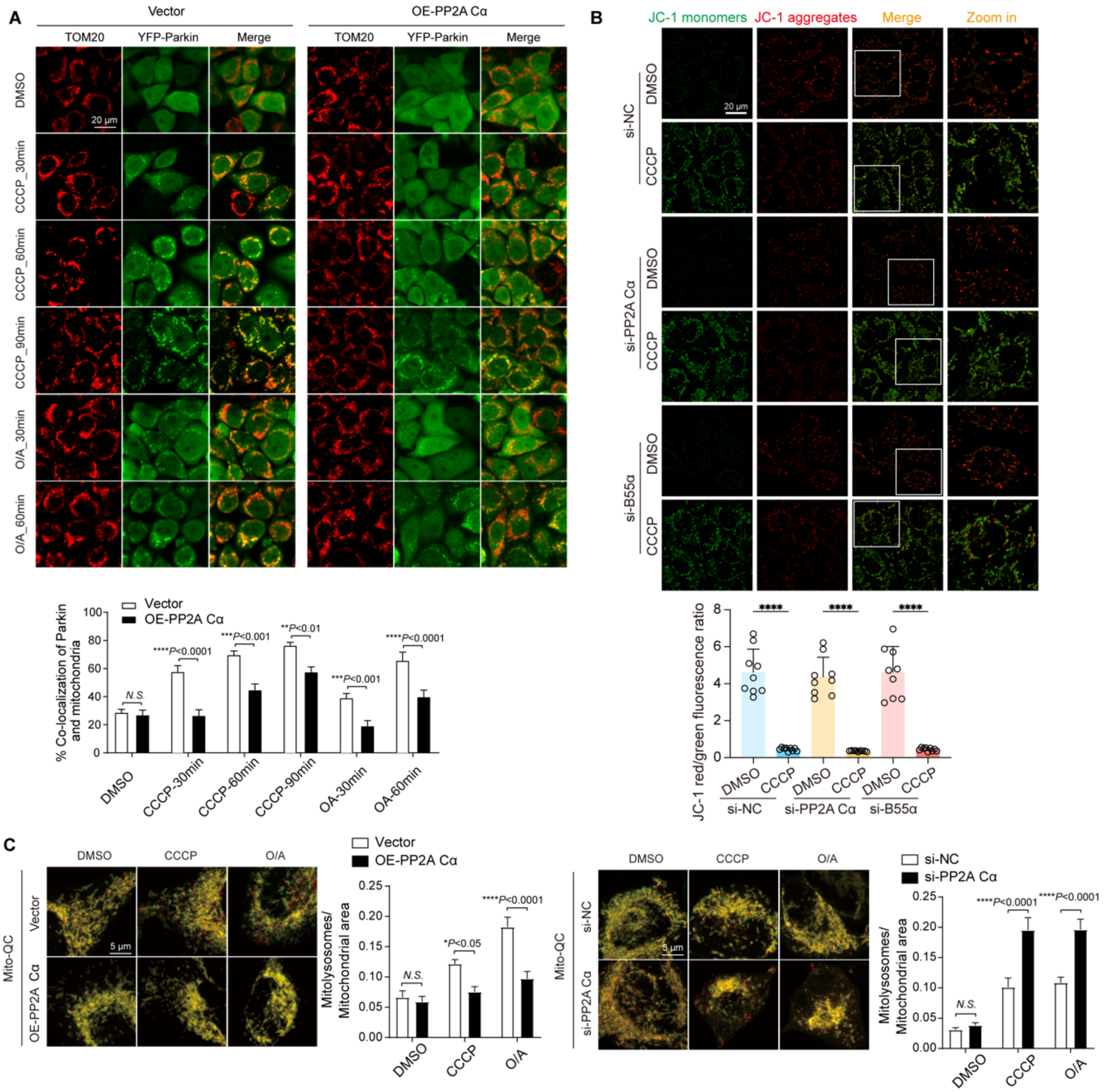


**Fig. S4. PP2A catalytic subunit modulates Parkin translocation and mitophagic flux. (A)** Time-course analysis of Parkin translocation. HeLa cells stably expressing YFP-Parkin (green) were transfected with an empty vector or untagged PP2A Cα, and treated with CCCP (10 μM) or O/A (250 nM) for the indicated durations. Mitochondria were stained for TOM20 (red). The graph below the representative images shows the quantitative analysis of Parkin colocalization with TOM20. (**B**) Assessment of mitochondrial membrane potential by JC-1 staining following PP2A Cα or B55α depletion. HeLa cells were transfected with non-targeting control siRNA (si-NC) or siRNA targeting PP2A Cα or B55α, followed by the treatment with DMSO or CCCP (10 μM for 1 h). JC-1 monomers and aggregates are shown in green and red, respectively. JC-1 only aggregates onto healthy mitochondria and emits red fluorescence. Boxed regions in the merged images are enlarged in the rightmost column. The graph below shows the JC-1 red/green fluorescence intensity ratio. No significant differences were detected among the siRNA groups under either DMSO- or CCCP-treated conditions. **(C)** PP2A modulates mitophagic flux as assessed by the Mito-QC reporter. SH-SY5Y cells expressing the Mito-QC reporter were transfected with Vector or PP2A Cα plasmid (left), or with si-NC or siRNA specific for PP2A Cα (right). Cells were treated with DMSO, CCCP (10 μM for 4 h), or O/A (both at 250 nM for 2 h). Mitophagy is indicated by the formation of red-only puncta (mito-lysosomes). The graphs beside the representative images show the quantitative analysis of the mito-lysosome area relative to the total mitochondrial area. Data in (A, C) are presented as mean ± SEM from n = 3 independent experiments. Data in (B) are presented as mean ± SEM across nine imaging fields per condition collected from 3 independent experiments; each dot represents one imaging field. Statistical significance was assessed by two-way ANOVA for (A, C) and by one-way ANOVA for (B). $^{*}P < 0.05$, $^{**}P < 0.01$, $^{***}P < 0.001$, and $^{****}P < 0.0001$; N.S., not significant.

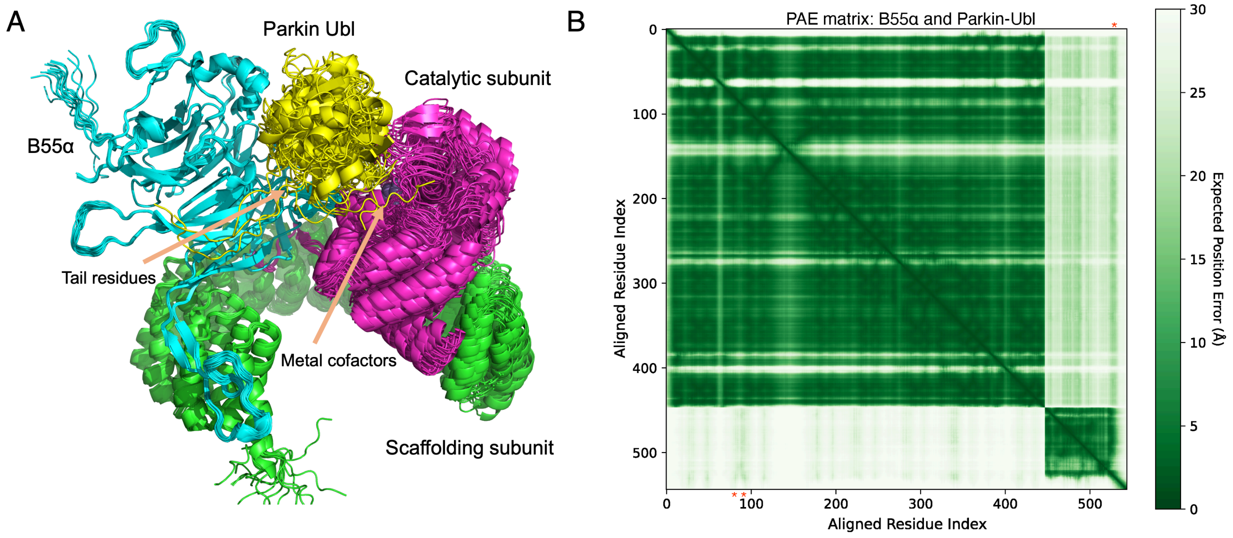


**Fig. S5. Structural convergence and validation metrics of the predicted PP2A–B55α–Parkin Ubl complex. (A)** Superposition of 15 converged, productive structural models (out of 25 total models generated across 5 independent AlphaFold 3 runs with different random seeds), superimposed onto the B55α subunit. Arrows highlight the basic C-terminal tail residues of the Ubl domain and the active-site binuclear metal cofactors in the catalytic subunit. While productive models were evaluated based on inter-subunit PAE values and the distance between Parkin pSer$^{65}$ and the binuclear metal center, rigid-body positional variations reflect intrinsic conformational flexibility. **(B)** Representative Predicted Aligned Error (PAE) matrix between B55α (residues 1–447) and the Parkin Ubl domain (residues 1–88; residue indices 448–535), taken from the prediction for the complex between heterotrimeric PP2A-B55α holoenzyme and Parkin Ubl domain (ipTM=0.76). Red asterisks denote the acidic β-hairpin on blade 4 of B55α (bottom left) and the basic C-terminal tail of the Ubl domain (top right) with low expected position errors. Representative atomic coordinates of the converged structural model are provided as Supplementary Data File S1.

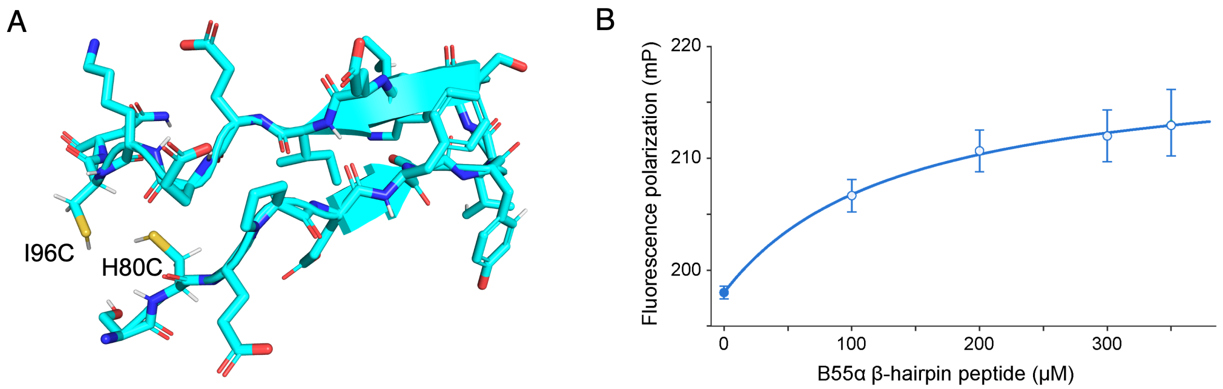


**Fig. S6. Direct *in vitro* binding between the B55α β-hairpin and Parkin Ubl assessed by fluorescence polarization. (A)** Structural model of the B55α β-hairpin peptide (residues 79–97 with native sequence of SHEPEFDYLKSLEIEEKIN), engineered with cysteine substitutions at positions 80 and 96 to stabilize the structure in solution. **(B)** Fluorescence polarization (FP) binding titration of FITC-labeled Parkin Ubl peptide (PP#1, FITC-QRPWRKG-CONH2, 5 μM in PBS buffer) titrated with increasing concentrations of the unlabeled, disulfide-constrained B55α peptide as shown in (A). Concentration-dependent binding saturation curve was fitted with on-site binding isotherm, affording a $K_d$ value of 133±108 μM. Data points represent mean ± SD from $n$ = 3 independent replicated FP measurements.

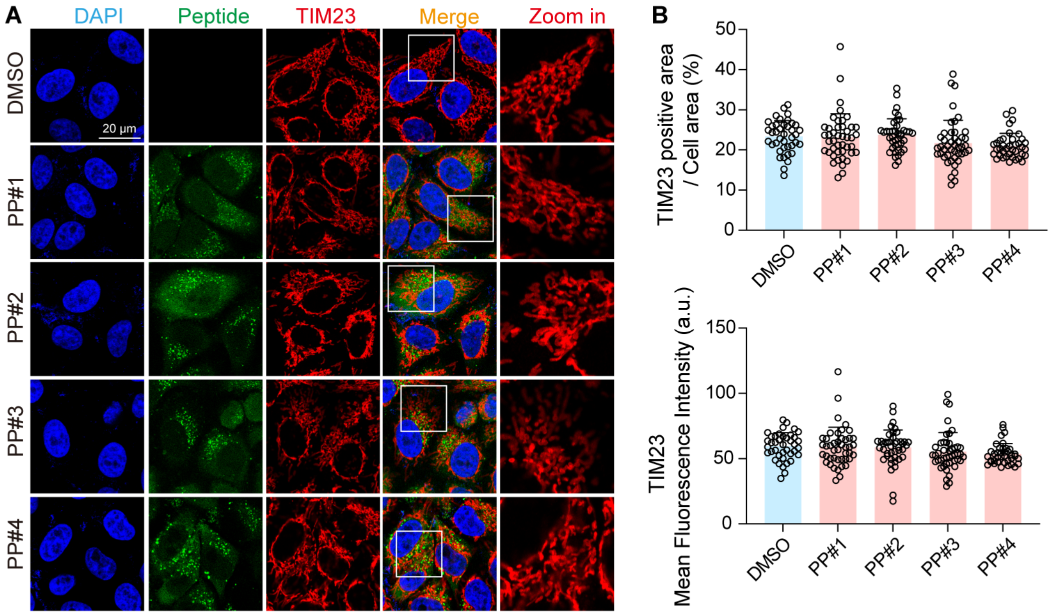


**Fig. S7 Effects of Ubl tail-derived peptides on mitochondrial morphology under basal conditions. (A)** Representative immunofluorescence images of cells treated with DMSO or the indicated FITC-labeled peptides (green) under basal conditions (without CCCP for mitochondria depolarization). Mitochondria were immunostained for TIM23 (red). **(B)** Quantitative analysis of the TIM23-positive area per cell area (%) and the mean fluorescence intensity of TIM23 from (A). Data are presented as mean ± SEM from $n$ = 3 independent experiments. No significant differences were observed among the groups, determined by one-way ANOVA.

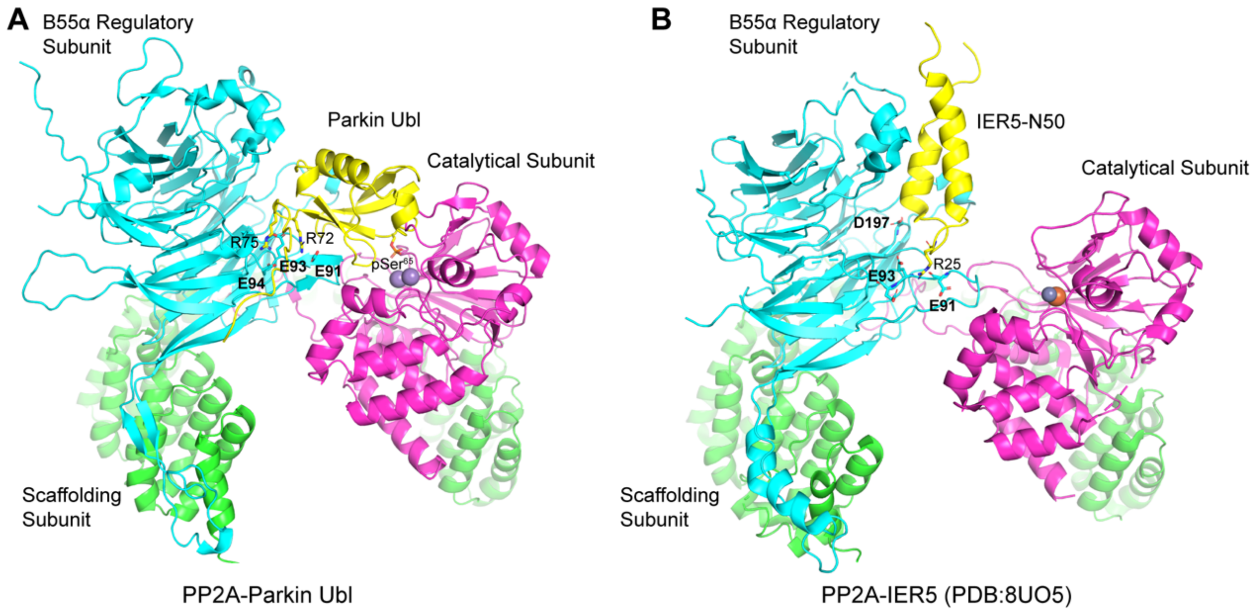


**Fig. S8. Comparison of the PP2A–B55α–Parkin Ubl complex structural model with the experimentally determined cryoEM structure of inhibitor-bound PP2A holoenzyme. (A)** AlphaFold model of the PP2A-B55α-Parkin Ubl complex. The scaffolding subunit is shown in green, the B55α regulatory subunit in cyan, the catalytic subunit in magenta, and the Parkin Ubl domain in yellow, with key interacting residues indicated. **(B)** For comparison, the cryo-EM structure of the PP2A-B55α holoenzyme bound to the specific inhibitor IER5 (PDB: 8UO5) is illustrated. IER5-N50 (yellow) engages both the canonical top groove (D197) and the lower β-hairpin interface via basic residues (e.g., R25 in IER5 with E91 and E93 in B55α), conceptually mirroring the non-canonical binding mode of the Parkin Ubl tail discovered in the present study.

**Table S1. Key resources table.**

| Reagent type (species) or resource | Designation | Source or reference | Identifiers | Additional information |
|---|---|---|---|---|
| Cell lines (Hs) | HeLa | CAS Center for Excellence in Molecular Cell Science, China | | |
| Cell lines (Hs) | YFP-Parkin-HeLa | PMID 19029340 | | A kind gift from Prof. Richard Youle |
| Cell lines (Hs) | SH-SY5Y | CAS Center for Excellence in Molecular Cell Science, China | | |
| Transfected construct (Hs) | PP2A-Cα-IRES-Aα | This paper | | See Protein sample preparation (Materials and Methods) |
| Transfected construct (Hs) | Parkin | This paper | | See Protein sample preparation (Materials and Methods) |
| Transfected construct (Hs) | PP2A-Cα-mCherry | This paper | | See Protein sample preparation (Materials and Methods) |
| Transfected construct (Hs) | Parkin-R72A | This paper | | See Cell Transfection and Treatment (Materials and Methods) |
| Transfected construct (Hs) | Parkin-R75A | This paper | | See Cell Transfection and Treatment (Materials and Methods) |
| Transfected construct (Hs) | Parkin-R72/75A | This paper | | See Cell Transfection and Treatment (Materials and Methods) |
| Transfected construct (Hs) | PP1α | This paper | | See Cell Transfection and Treatment (Materials and Methods) |
| Transfected construct (Hs) | B55α-3xFlag | This paper | | See Cell Transfection and Treatment (Materials and Methods) |

| | | | | |
|---|---|---|---|---|
| Transfected construct (Hs) | B55α (F84A)-3xFlag | This paper | | See Cell Transfection and Treatment (Materials and Methods) |
| Transfected construct (Hs) | B55α (E91A)-3xFlag | This paper | | See Cell Transfection and Treatment (Materials and Methods) |
| Transfected construct (Hs) | B55α (E93A)-3xFlag | This paper | | See Cell Transfection and Treatment (Materials and Methods) |
| Transfected construct (Hs) | B55α (E94A)-3xFlag | This paper | | See Cell Transfection and Treatment (Materials and Methods) |
| Transfected construct (Hs) | B55α (E91/E93/E94A)-3xFlag | This paper | | See Cell Transfection and Treatment (Materials and Methods) |
| Transfected construct (Hs) | siRNA to PP2A Cα | This paper | | See Cell Transfection and Treatment (Materials and Methods) |
| Transfected construct (Hs) | siRNA to PP2A B55α | This paper | | See Cell Transfection and Treatment (Materials and Methods) |
| Transfected construct (Hs) | Control siRNA | This paper | | See Cell Transfection and Treatment (Materials and Methods) |
| Antibody | Anti-Parkin | Cell Signaling Technology | Cat# 2132S | WB (1:1000) |
| Antibody | Anti-phospho-Parkin (Ser65) | Cell Signaling Technology | Cat# 36866S | WB (1:500) |
| Antibody | Anti-ubiquitin | Beyotime Biotechnology | Cat# AF1705 | WB (1:1000) |
| Antibody | Anti-phospho-ubiquitin (Ser65) | Merck Millipore | Cat# ABS1513-I | WB (1:500) |
| Antibody | Anti-B55α | Affinity Biosciences | Cat# DF6908 | WB (1:1000) |
| Antibody | Anti-PP2A Cα | Invitrogen | Cat# MA5-32920 | WB (1:2000) |
| Antibody | Anti-TIM23 | Proteintech | Cat# 11123-1-AP | WB (1:5000) |
| Antibody | Anti- PP1α | Proteintech | Cat# 85445-3-RR | WB (1:5000) |
| Antibody | Anti-PPP2R3A | Genetex | Cat# 11123-1-AP | WB (1:1000) |

| | | | | |
|---|---|---|---|---|
| Antibody | Anti-GAPDH | Proteintech | Cat# 60004-1-Ig | WB (1:20000) |
| Antibody | HRP-conjugated goat anti-rabbit IgG | Beyotime Biotechnology | Cat# A0208 | WB (1:2000) |
| Antibody | HRP-conjugated goat anti-mouse IgG | Beyotime Biotechnology | Cat# A0216 | WB (1:2000) |
| Antibody | Anti-TOM20 | Santa Cruz | Cat# SC-17764 | IF (1:200) |
| Antibody | Anti-TIM23 | Proteintech | Cat# 11123-1-AP | IF (1:200) |
| Antibody | Anti-Parkin | CST | Cat# 4211 | IF (1:100) |
| Antibody | Anti-Goat Mouse IgG (H+L) cross-adsorbed secondary antibody with Alexa Fluor 647 | Thermo Fisher Scientific | Cat# A-21235 | IF (1:200) |
| Antibody | Anti-Goat Rabbit IgG (H+L) cross-adsorbed secondary antibody with Alexa Fluor 594 | Thermo Fisher Scientific | Cat# A-11080 | IF (1:200) |
| Recombinant protein (*Hs*) | Ub | PMID 28611216 | | |
| Recombinant protein (*Hs*) | K48C Ub | This paper | | See Protein sample preparation (Materials and Methods) |
| Recombinant protein (*Hs*) | T66V/L67N Ub | PMID 29133469 | | |
| Recombinant protein (*Hs*) | FLAG-Ub | This paper | | See Protein sample preparation (Materials and Methods) |
| Recombinant protein (Hs) | His-PP1α | This paper | | See Protein sample preparation (Materials and Methods) |
| Recombinant protein (*Hs*) | His-PP2A Cα | This paper | | See Protein sample preparation (Materials and Methods) |
| Recombinant protein (*Hs*) | GST-PP2A Aα | This paper | | See Protein sample preparation (Materials and Methods) |
| Recombinant protein (*Hs*) | GST-B55α | This paper | | See Protein sample preparation |

| | | | | |
|---|---|---|---|---|
| | | | | (Materials and Methods) |
| Recombinant protein (*Pediculus humanus*) | GST-*Ph*PINK1 (115-575) | PMID 28611216 | | |
| Recombinant protein (*Hs*) | GST-Ubl (1-76) | This paper | | See Protein sample preparation (Materials and Methods) |
| Recombinant protein (*Hs*) | His-SUMO-Parkin | This paper | | See Protein sample preparation (Materials and Methods) |
| Peptide | Poly-FLAG peptide | Bimake | Cat# B23111 | |
| Commercial assay or kit | Phos-tag acrylamide | Wako Chemicals | Cat# AAL-107 | |
| Commercial assay or kit | BCA protein assay kit | Beyotime Biotechnology | Cat# P0012S | |
| Commercial assay or kit | RIPA buffer | Beyotime Biotechnology | Cat# P0013C | |
| Commercial assay or kit | Protease inhibitors | Coolaber | Cat# SL1086 | |
| Commercial assay or kit | Phosphatase inhibitors | Coolaber | Cat# SL1087 | |
| Commercial assay or kit | $BS^3$ | Thermo Fisher | Cat# 21580 | |
| Commercial assay or kit | Anti-Flag magnetic beads | Bimake | Cat# B26102 | |
| Commercial assay or kit | Clarity™ Western ECL Substrate | Bio-Rad Laboratories | Cat# 1705060 | |
| Commercial assay or kit | Normal donkey serum | Yeasen Biotechnology | Cat# 36116ES03 | |
| Commercial assay or kit | JC-1 | Beyotime | Cat# C2006 | |
| Commercial assay or kit | DAPI Fluoromount-G™ | Yeasen Biotechnology | Cat# 36308ES11 | |
| Chemical compound | 3-Bromo-1,1,1-trifluoroacetone | J&K Scientific | Cat# 312226 | CAS No. 431-35-6 |
| Chemical compound | Okadaic Acid | Aladdin | Cat# O275664 | CAS No. 209266-80-8 |
| Chemical compound | Cantharidin | Aladdin | Cat# C111020 | CAS No. 56-25-7 |
| Chemical compound | CCCP | Sigma-Aldrich | Cat# C2759 | CAS No. 555-60-2 |
| Chemical compound | Oligomycin | Abcam | Cat# ab141829 | CAS No. 1404-19-9 |
| Chemical compound | Antimycin A | Sigma-Aldrich | Cat# A8674 | CAS No. 1397-94-0 |
| Software, algorithm | pFind | PMID 30295672 | | |
| Software, algorithm | NMRPipe | PMID 8520220 | | |

| | | |
|---|---|---|
| Software, algorithm | NMRViewJ | PMID 22911360 |
| Software, algorithm | TopSpin | Bruker |
| Software, algorithm | Origin | OriginLab |
| Software, algorithm | Image Lab | Bio-Rad Laboratories |
| Software, algorithm | ImagJ | NIH |
| Software, algorithm | PyMOL | Schrodinger |
| Software, algorithm | NIS-Elements AR | Nikon |
| Software, algorithm | Prism | GraphPad |
